# Schwann Cell Mapping and Characterization in Bone of Different Embryonic Origins

**DOI:** 10.64898/2026.05.28.727991

**Authors:** Anushka Gerald, Quentin Meslier, Mohamed G Hassan, Isabella Rastegar, Erica L Scheller

## Abstract

Schwann cells (SCs) provide support for nerves throughout the body. Despite importance for nerve function and repair, the morphology and distribution of SCs in bone remains largely undefined. In this study we used a “Schwann Cell Mapper” mouse (*Mpz*-Cre^+/-^;TdT^+/+^;*Ngfr*-eGFP^+/+^) and a “p75 Lineage Tracer” mouse (*Ngfr*-CreERT2^+/-^;ZsGreen1^+/+^) to study SC localization and morphology within the adult mouse calvaria, limb, and vertebrae. We found that all nerves in bone were covered by mature myelinating or non-myelinating SCs, labeled by MPZ and p75-NGFR, respectively. Mature SCs populated the periosteum and entered the bone marrow through transcortical canals. Non-myelinating SCs outnumbered myelinating SCs in bone, with a ratio of ∼2:1 by length density. Non-myelinating SCs in periosteum had more branching and increased size relative to myelinating SCs. In addition, we identified two candidate populations of MPZ lineage+ and p75-NGFR+ immature SCs (iSCs) that were distributed throughout the calvarial periosteum. Similar to mature SCs, p75-NGFR+ candidate iSCs were more prevalent at a ratio of ∼3:1. Overall, neural crest-derived calvarial bone had evidence of increased SC maturity relative to mesoderm-derived sites, identifying niche-level differences in SC maturation. Lastly, lineage tracing revealed that mesenchymal lineages in bone were largely negative for both MPZ and p75-NGFR (<0.1% labeling). These findings provide a framework of SC organization in bone and highlight previously unrecognized diversity across skeletal compartments. By defining distribution and morphology, this work lays the foundation for future studies investigating how SCs contribute to bone biology, including roles in repair, pain, and homeostasis.

## INTRODUCTION

Schwann cells (SCs) provide trophic and structural support for nerve axons throughout the body and have unique plasticity allowing for transition into a repair state after local injury. Emerging evidence suggests that this is also true for bone. Specifically, five *in vivo* reports show that SC lineages can produce secretory factors including platelet-derived growth factor-AA (PDGF-AA) and oncostatin M (OSM) that accelerate bone repair.(Johnston et al., 2016; Jones et al., 2019; Zhang et al., 2023; Shen et al., 2025; Yang et al., 2026) *In vitro*, SCs enhance endothelial cell migration, promote proliferation and differentiation of osteoprogenitors, and inhibit osteoclastogenesis.(Cai et al., 2010; Wang et al., 2025, 2026; Yang et al., 2026) In the bone marrow, non-myelinating SCs are thought to maintain the hematopoietic niche by regulating activation of transforming growth factor-beta (TGF-β).(Yamazaki et al., 2011) The regenerative capacity of SCs has been leveraged to improve bone healing through methods such as direct SC transplantation, delivery of SC conditioned media, or introduction of isolated SC exosomes to the injury site.(Wu et al., 2020; Zhang et al., 2021; Hao et al., 2023; Wang et al., 2023, 2025; Cui et al., 2025) SC function has also been linked to bone pain.(Landini et al., 2023) Despite the emerging importance of SCs for nerve function and tissue repair in bone, the morphology and distribution of myelinating and non-myelinating SCs in skeletal tissues remains largely undefined.

Mature SCs derive from the neural crest. In mice, this begins at ∼E13 in conjunction with the finalization of bone patterning, whereby neural crest stem cells (NCSCs) transition into SC precursors (SCPs).(Wanner et al., 2006; Jessen et al., 2015) SCPs connect to each other with sheet-like processes and adherens junctions to start enveloping axons.(Wanner et al., 2006) Compared to NCSCs, SCPs are nerve associated, express glial differentiation factors, and are dependent on axon-associated survival signal neuregulin 1 type III.(Dong et al., 1999; Buchstaller et al., 2004; Jessen and Mirsky, 2005) SCPs provide trophic support of peripheral axons and are important for nerve fasciculation.(Jessen and Mirsky, 2005) SCPs mostly differentiate into immature Schwann cells (iSCs) but can also become fibroblast, osteoblast, and chondrocyte lineages *in vitro*.(Joseph et al., 2004) Despite this potential, lineage tracing in mice *in vivo* reveals that only 0.5 to 2% of embryonic OSX+ osteolineage cells derive from SCPs in the mandible, rib, and scapula.(Xie et al., 2019) Furthermore, this only occurs during a limited developmental window during E12.5-E17.5.(Xie et al., 2019) In later embryonic stages in mice and in adult fish, SCPs tagged during early development can give rise to osteocytes and clonal populations of chondrocytes.(Xie et al., 2019) However, this phenomenon is rare. The contributions of SCPs to *in vivo* osteolineages in adult mice remains unknown.

Unlike SCs, bone can arise from multiple embryonic origins. The calvaria provides an effective model to study the differences between adjacent neural crest-and mesoderm-derived bone. For example, lineage tracing studies have shown that the frontal bone is neural crest in origin whereas the nearby parietal bone is mesoderm in origin.(Chai et al., 2000; Jiang et al., 2002) This can be contrasted with other mesoderm derived bones in the rest of the skeleton.

Previous literature in other organ systems pinpoints myelin protein zero (MPZ) and p75 nerve growth factor receptor (p75-NGFR) as markers of myelinating and non-myelinating SCs, respectively. (Yasuda et al., 1987; Bentley and Lee, 2000) In this study, we used two mouse models to determine if these markers could selectively label SCs in bone. Model #1 “Schwann Cell Mapper” is a triple mutant reporter mouse (*Mpz*-Cre^+/-^;TdT^+/+^;*Ngfr*-eGFP^+/+^) that expresses eGFP in cells that are actively expressing p75-NGFR and TdTomato in cells that previously or currently express MPZ-Cre. This model is intended to visualize myelinating (MPZ lineage+) and non-myelinating (p75-NGFR+) SCs simultaneously. It is also sufficient to define any skeletal cells arising from the MPZ+ lineage.

Model #2 “p75 Lineage Tracer” is a new CRISPR-generated, inducible p75-NGFR-CreERT2 line that expresses ZsGreen1 only after treatment with tamoxifen (*Ngfr*-CreERT2^+/-^;ZsGreen1^+/+^). This model is intended to label p75-NGFR+ populations including non-myelinating SCs. In this research, we validate MPZ and p75-NGFR as skeletal SC markers to visualize the intricate network of myelinating and non-myelinating SCs in adult bone of multiple embryonic origins, including frontal bone, parietal bone, tibia, femur, and vertebrae. We also confirm that SCs are present in the mouse calvaria at birth, identify unique patterns of myelinating vs non-myelinating SCs in bone, and, for the first time, localize candidate iSCs within the periosteum.

## MATERIALS AND METHODS

### Animals

All work was performed as approved by the animal use and care committee at Washington University (St. Louis, MO, USA). Mice were housed on a 12-hour light/dark cycle and fed ad libitum (PicoLab 5053, LabDiet, St. Louis, MO, USA). SC mapper mice (*Mpz*-Cre^+/-^;TdT^+/+^;*Ngfr*-eGFP^+/+^) were a kind gift of Dr. Jeffrey Milbrandt and were generated by crossing the following strains: MPZ-Cre (JAX Strain 017927)(Feltri et al., 1999), Ai9 (JAX Strain 007909, producing the TdTomato reporter) and p75NGFR-Cre (Mutant Mouse Resource & Research Center, strain 036182-UCD).(Gong et al., 2003) P75 Lineage Tracer mice (*Ngfr*-CreERT2^+/-^;ZsGreen1^+/+^) were generated by crossing new, CRISPR-generated p75-NGFR-CreERT2 mice with Ai6 (JAX Strain 007909, producing the ZsGreen reporter). The p75-NGFR-CreERT2 model was created by the Genome Engineering & Stem Cell Center (GESC@MGI) at Washington University as follows. Briefly, a synthetic gRNA (sgRNA) targeting the sequence, 5’-gtcgccggaggcgagcaatgagg was purchased from IDT (Coraville, IA) and validated for cleavage activities in Neuro 2a cells. A homology-directed repair donor with Cre-ERT2-P2A sequences flanked by 800 bp homology arms was cloned in between a pair of AAV2 inverted terminal repeats by Genscript (Nanjing, China), and recombinant AAV6 virus was prepared by the Hope Center Viral Vector Core at Washington University in St. Louis. The rAAV donor was co-electroporated with the sgRNA/Cas9 complex into Neuro2a cells to confirm integration at the N-terminus of the Ngfr coding sequence. Mouse production using the above validated reagents was done at the Mouse Genetics Core at Washington University in St. Louis and genotyped as described previously.(Sentmanat et al., 2026)

### Calvaria and soft tissue isolation, immunostaining and imaging

Mice were anesthetized with isofluorane and perfused through the left ventricle of the heart with 10 mL phosphate buffered saline followed by 10 mL 10% neutral buffered formalin (NBF; Thermo Fisher Scientific, Waltham, MA, USA; 23–245684). Calvaria were isolated by decapitating mice with a blade. Skin was gently degloved from the skull and removed. Curved scissors were used to cut around the margins of the calvaria, isolating it from the rest of the skull and brain. Soft tissues including heart, liver, intestine, kidney, pancreas, and spleen were also collected. Tissues were post-fixed in 10% NBF for 24-hours and then washed for 2 hours in diH_2_O. Calvaria were also fully decalcified in 14% EDTA, pH 7.2-7.4, over 5 days (Sigma-Aldrich, St. Louis, MO, USA; E5134).

Whole calvarial preparations were blocked and permeabilized in 10% normal donkey serum (NDS) and 0.3% Triton X-100 in PBS for 24 hours before incubation for 5 days with primary antibodies (1:1000 dilution in TNT +1% NDS (Supplemental Table 1) at 4°C. For, SC Mapper mouse samples, eGFP reporter was amplified using GFP primary antibodies (1:1000 dilution in TNT +1% NDS. After washing 2×15 minutes and 1x overnight, secondary antibodies (1:1000 dilution in TNT+1% NDS) were applied for 5 days at 4°C. The sections were then washed 3×15 minutes in TNT buffer, incubated in DAPI (Sigma-Aldrich) for 5 minutes, and stored in PBS until imaging. Calvaria were placed on a 35mm glass bottom dish with Fluoromount and covered with a glass coverslip. The coverslip was weighed down with a metal ring to flatten the sample. For Figs.1,2,4,5,7,8,9,10, serial tiled images were taken with a 10x, 20x, or 40x objective on a Nikon (Tokyo, Japan) W1 CSU SoRa Spinning Disk Confocal and Superresolution (μm/px = 0.650, step size = 10 μm) with automatic stitching.

For soft tissues, samples were cryoprotected using 30% sucrose in PBS for 24 hours at 4°C. Samples were embedded in OCT using a cold plate. Samples were then sectioned with a cryostat in 20µm sections and placed on colorfrost microscope slides. Slides were air-dried for 30 minutes and washed 2×5 min with 1x PBS. Slides were blocked and permeabilized with a 10% NDS and 0.3%

Triton X-100 in PBS for 1 hr. at room temperature (RT) before incubation for overnight with primary antibodies (1:1000 dilution in TNT +1% NDS (Supplemental Table 1) at RT in a humidified chamber. After washing 3×5 minutes, secondary antibodies (1:1000 dilution in TNT +1% NDS) were applied for 24 hours at 4°C. The sections were then washed 3×5 minutes in TNT buffer, and incubated in DAPI (Sigma-Aldrich) for 5 minutes. Slides were then coverslipped (Fisher Scientific 12-545-M) with Fluoromount-G (Thermo Fisher Scientific 00-4958-02), air dried, sealed with clear nail polish, and stored until imaging.

### Limb and vertebrae bone tissue clearing and light-sheet microscopy

For analysis of limb bones and vertebrae, mice were perfused with 10 mL of 1x PBS and 10 mL of 4% Paraformaldehyde. Following perfusion, hindlimb and caudal vertebrae were harvested and surrounding soft tissues were trimmed, taking care to preserve the integrity of the periosteum.

Samples were fixed in 4% paraformaldehyde at 4°C overnight with gentle agitation. After fixation, tissues were rinsed and decalcified in 14% EDTA, pH 7.2-7.4 for 10 days at 4°C, with the solution refreshed every 3 days. Following decalcification, samples were washed and processed using a solvent-based tissue clearing protocol.(Qi et al., 2019; Meslier et al., 2024) Briefly, samples were dehydrated in a graded series of tetrahydrofuran (THF; 50%, 70%, 80%, 95%, 95%) prepared in PBSQ (PBS containing 25% Quadrol), with 1-hour incubation at each step. Samples were then incubated overnight in dichloromethane for delipidation. The following day, tissues were rehydrated through a reverse THF gradient (95%, 95%, 80%, 70%, 50%) and washed extensively in PBS (five washes throughout the day). For immunolabeling, samples were incubated overnight in blocking buffer consisting of TNT buffer supplemented with 5% NDS. Tissues were then incubated with primary antibodies (1:400 dilution in TNT + 5% NDS, Supplemental Table 1) for 5 days at RT. After primary incubation, samples were washed three times for 1 hour in TNT buffer, followed by an overnight wash. Secondary antibodies (1:250 dilution in TNT + 5% NDS) were applied under the same conditions, followed by identical washing steps. Nuclear staining was performed by overnight incubation with DAPI in TNT buffer (1:750). Finally, samples were subjected to refractive index matching using EasyIndex (RI = 1.52) overnight and subsequently imaged using Miltenyi Ultra Microscope Blaze (4X/0.35, and 12X/0.53 objectives, XY resolution: 1.62 µm and 0.54 µm respectively, Z resolution: 2µm).

### Image processing and quantification

For confocal datasets, the.nd2 images were converted to.ims files for processing in Imaris (Oxford Instruments, Version 11). P75-NGFR+ and MPZ+ SCs were manually traced in 3D to create filaments in Imaris. SC filaments were identified by local contrast with the background, clear linear structures, and bright nuclei along the filament. These filaments were then converted into surfaces. The surface area of the bone was calculated based on total image area. Density of fiber lengths was calculated as the total length of fibers traced divided by the corresponding bone surface area for the region traced. The estimated length of SCs was calculated by dividing the total length of the SC filaments by the number of nuclei along the SC filaments. P75-NGFR+ and MPZ+ iSCs were identified morphologically and by their bright large nuclei using Imaris’s spot function. SC branching was quantified by manually counting the number of processes extending directly from the nucleus.

Branching events occurring distal to the nucleus were not included. files for visualization in Imaris (Oxford Instruments, Version 11). Maximum projection views were generated using the Imaris ortho slicer visualization tool.

### Statistics

Statistical comparisons were performed in GraphPad Prism. Values compared within multiple biological sites in one mouse or between two reporters in mouse were evaluated by a paired t-test. Values compared between two independent groups were evaluated using a Welch’s-test due to unequal sample sizes. Individual data points are presented in the figures and represent biological replicates (individual mice).

## RESULTS

### MPZ and p75-NGFR label nerve-lining Schwann cells in skeletal tissues

MPZ^+^ myelinating SCs are known to cover single larger axons, providing wraps of myelin to support rapid nerve conduction.(Peters and Muir, 1959; Rasminsky and Sears, 1972; Lodin et al., 1973) By contrast, p75-NGFR+ non-myelinating SCs can surround and support multiple small axons simultaneously in structures termed Remak bundles.(Remak, 1838; Rasminsky and Sears, 1972) Whole mount images of the calvaria from adult male and female ‘Schwann Cell Mapper’ mice at 12 to 24 weeks of age demonstrated broad distribution of MPZ lineage^+^ and p75-NGFR^+^ cells overlying TUBB3^+^ nerve axons throughout the calvarial periosteum (Fig.1A). Upon close examination, it appeared that all TUBB3^+^ axons had an associated MPZ lineage or p75-NGFR^+^ cell along its entire length, though the signal was sometimes faint (Fig.1B). Nerve-associated MPZ lineage^+^ and p75-NGFR^+^ cells co-stained for S100B, a marker of SCs (Fig.2A). Comparable net-like patterns of periosteal p75-NGFR^+^ nerve-associated cells were observed in the ‘p75 Lineage Tracer’ model 48-hours after labeling with tamoxifen in male and female mice at 14 to 21 weeks of age (Fig.1B,2A).

**Figure 1.**
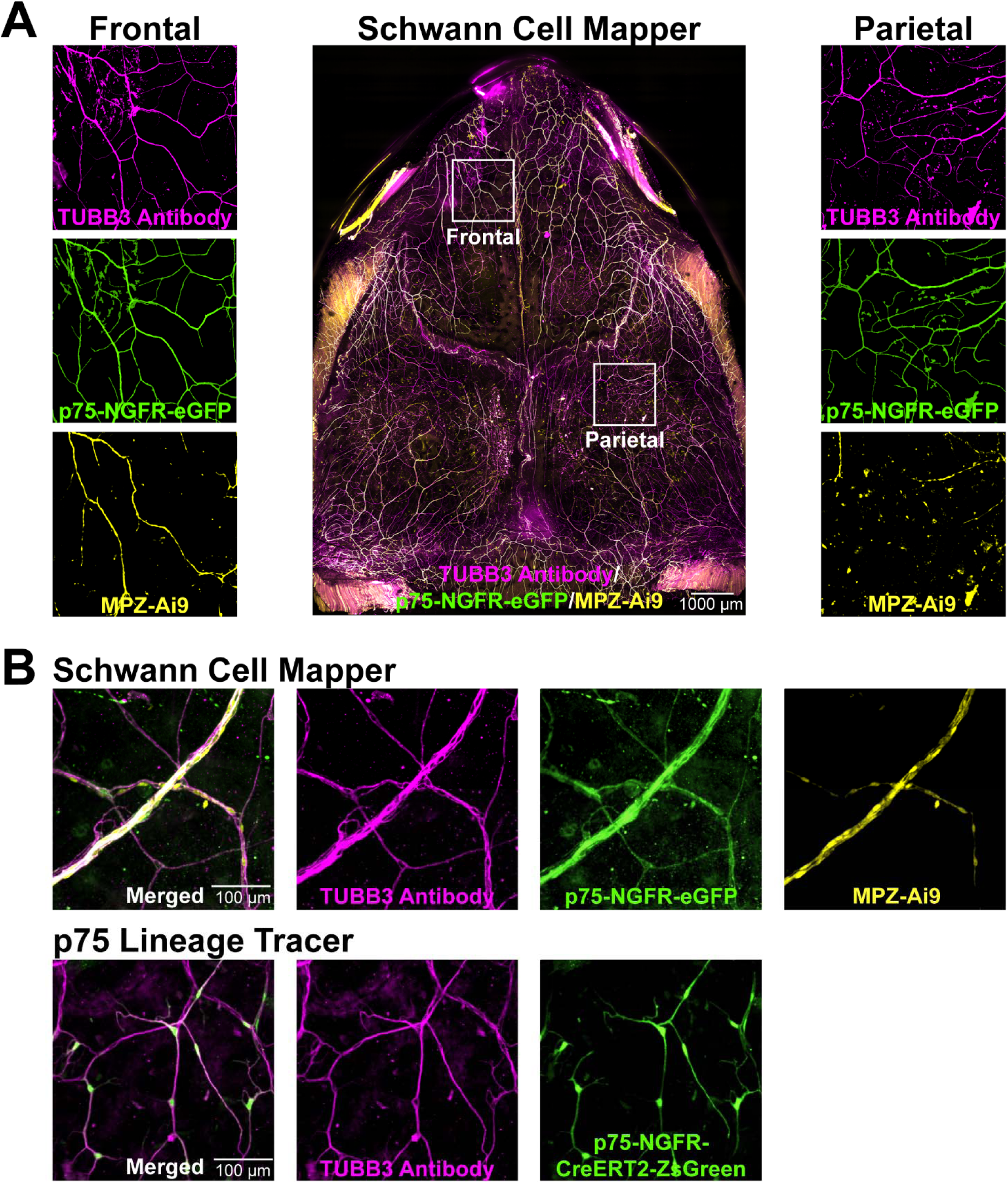
Distribution of MPZ lineage and p75-NGFR+ Schwann cells in the calvaria and association with peripheral nerves. **(A)** Representative maximum intensity projection (MIP) of whole mount adult ‘Schwann Cell Mapper’ mouse calvaria (*Mpz*-Cre^+/-^;TdT^+/+^;*Ngfr*-eGFP^+/+^), imaged at 4x resolution on a Nikon spinning disk confocal. White boxes represent zoomed in regions displayed to the left (frontal bone) and right (parietal bone). Scale = 1000 µm. **(B)** MIP of whole mount 10x spinning disk confocal images of adult ‘Schwann Cell Mapper’ (top) and ‘p75 Lineage Tracer’ (bottom, *Ngfr*-CreERT2^+/-^;ZsGreen1^+/+^) mouse calvarial periosteum showing colocalization with TUBB3+ nerve axons. Scale = 100 µm. Magenta = TUBB3 antibody for nerve axons, Green = p75-NGFR-eGFP or p75-NGFR-CreERT2-ZsGreen reporter for non-myelinating SCs, Yellow = MPZ-Ai9 reporter for myelinating SCs.

**Figure 2.**
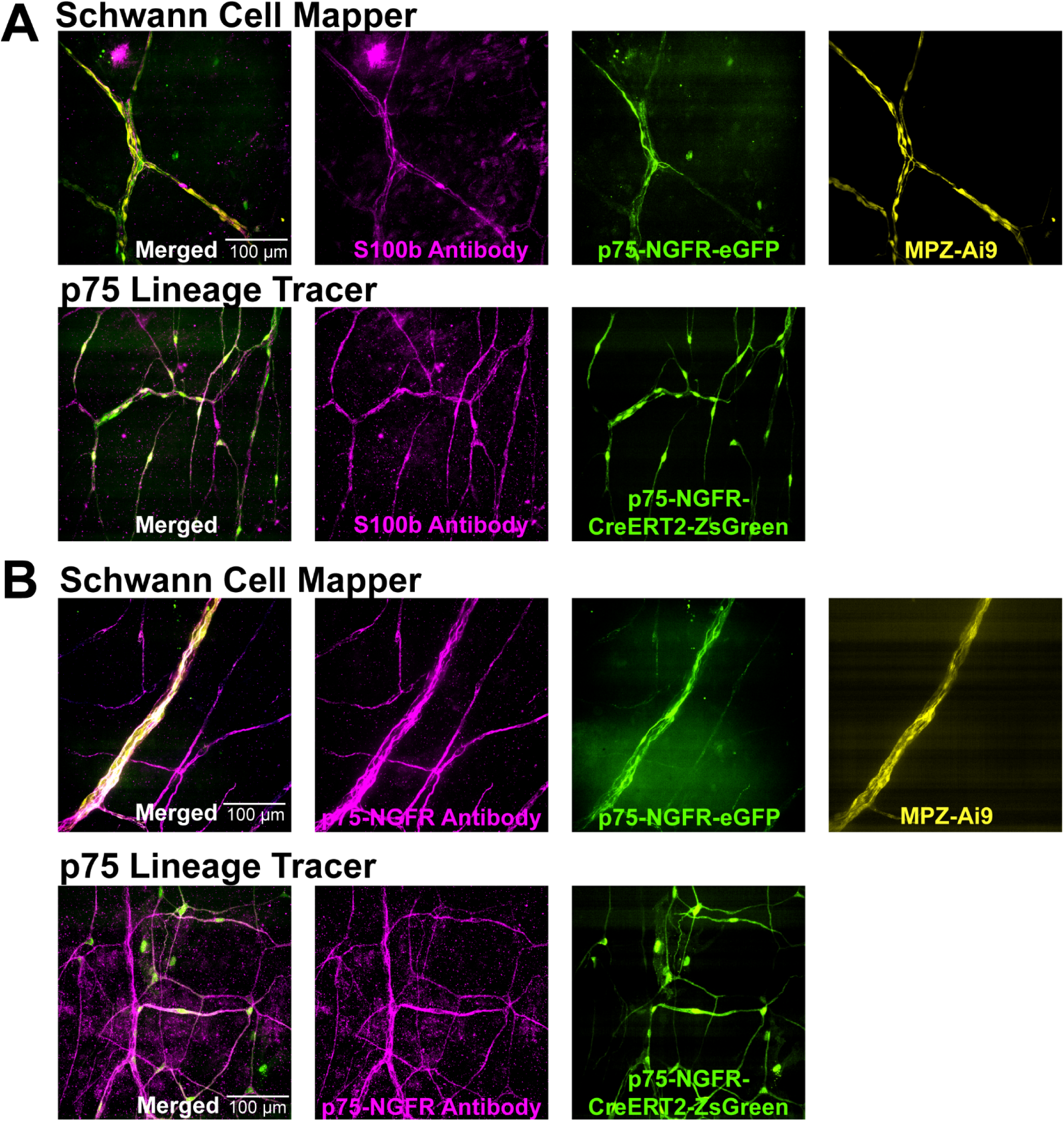
MPZ and p75-NGFR Schwann cell reporter co-staining with antibodies against S100B and p75-NGFR. **(A)** Maximum intensity projection of whole mount 10x spinning disk confocal images of adult ‘Schwann Cell Mapper’ (top, *Mpz*-Cre^+/-^;TdT^+/+^;*Ngfr*-eGFP^+/+^) and ‘p75 Lineage Tracer’ (bottom, P75-CreERT2^+/-^;ZsGreen1^+/+^) mouse calvarial periosteum showing colocalization with (A) S100B antibody and (B) p75-NGFR antibody. Scale = 100 µm. Magenta = S100B or p75-NGFR antibody, as indicated, Green = p75-NGFR-eGFP or p75-NGFR-CreERT2-ZsGreen reporter for non-myelinating SCs, Yellow = MPZ-Ai9 reporter for myelinating SCs.

These cells also co-stained with antibodies targeting S100B (Fig.2A) and p75-NGFR (Fig.2B).

Similar distributions of nerve-lining MPZ lineage^+^ and p75-NGFR^+^ cells were observed within the tibia, as visualized with tissue clearing and lightsheet imaging of ‘Schwann Cell Mapper’ mice (Fig.3). In long bone, MPZ lineage^+^ nerve-lining cells were concentrated within the outer fibrous layer of the periosteum and p75-NGFR^+^ cells within the cambium (Supplemental Fig.1). Both MPZ lineage^+^ and p75-NGFR^+^ cell types had central, rounded nuclei and long, thin cytoplasmic extensions, overlaying TUBB3^+^ nerves with a “bead on a string” morphology (Fig.1,2). Based on reporter expression, morphology, and co-stain with S100B, these findings are consistent with nerve-associated MPZ lineage^+^ and p75-NGFR^+^ cells being mature SCs.

**Figure 3.**
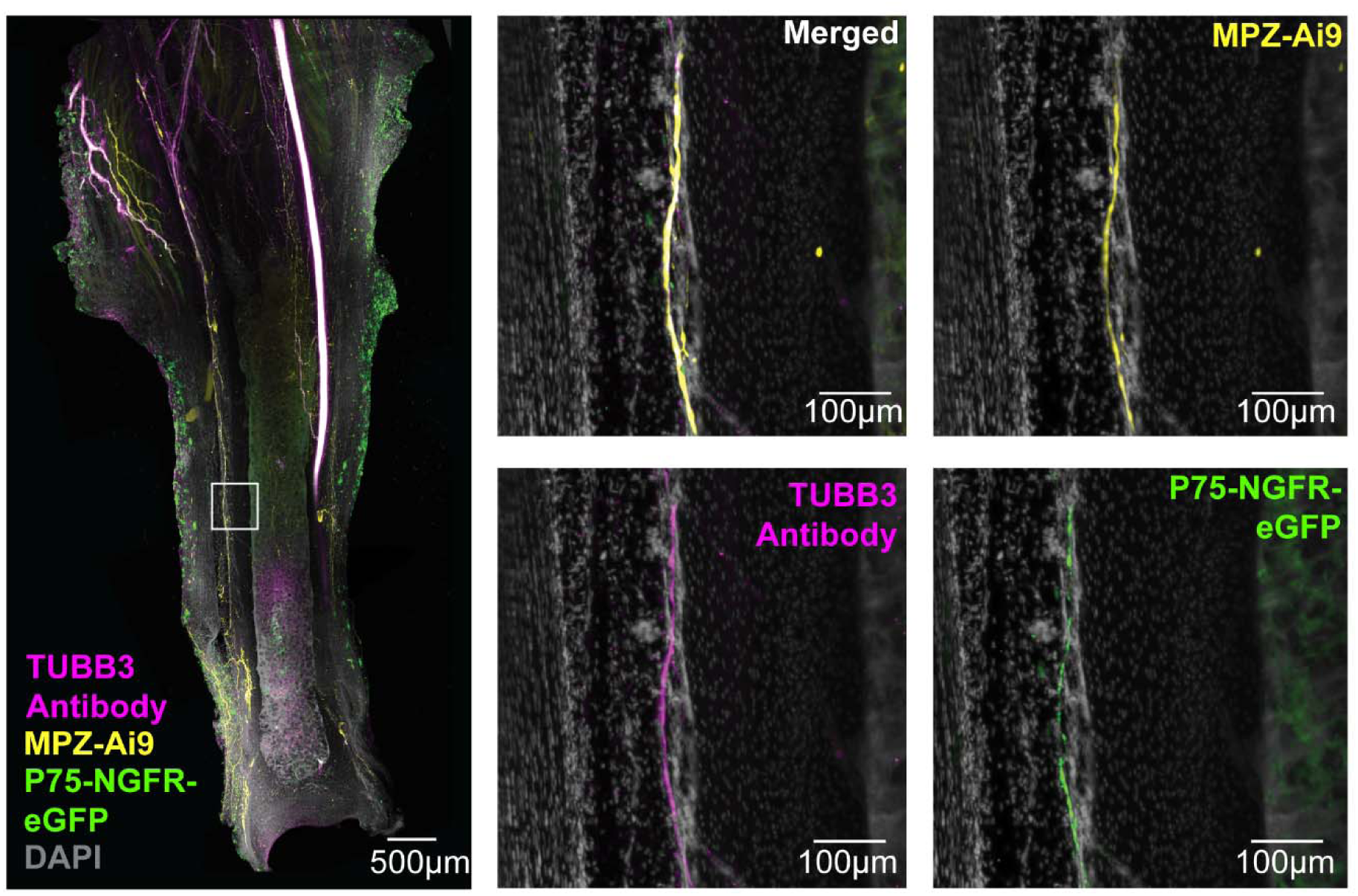
Schwann cells and nerve in mouse distal tibia. Maximum intensity projection of whole mount 4x lightsheet microscopy images of a ‘Schwann Cell Mapper’ (*Mpz*-Cre^+/-^;TdT^+/+^;*Ngfr*-eGFP^+/+^) mouse showing presence of nerves, myelinating SCs, and non-myelinating SCs in mouse tibia periosteum. Scale = 500µm. The white box represents a zoomed in region of the distal tibia. Scale= 100 µm. Magenta = TUBB3 antibody, Yellow = MPZ-Ai9 reporter for myelinating SCs, Green = p75-eGFP reporter for non-myelinating SCs, Gray = DAPI.

### Periosteal MPZ and p75-NGFR Schwann cells are present at birth

To determine if MPZ lineage^+^ and p75-NGFR^+^ SCs within the periosteum were already present at birth, we imaged whole mount calvaria from ‘Schwann Cell Mapper’ mice at postnatal day 0. As in adult mice, both MPZ lineage^+^ and p75-NGFR^+^ SCs were prevalent within the calvarial periosteum (Fig.4).

**Figure 4.**
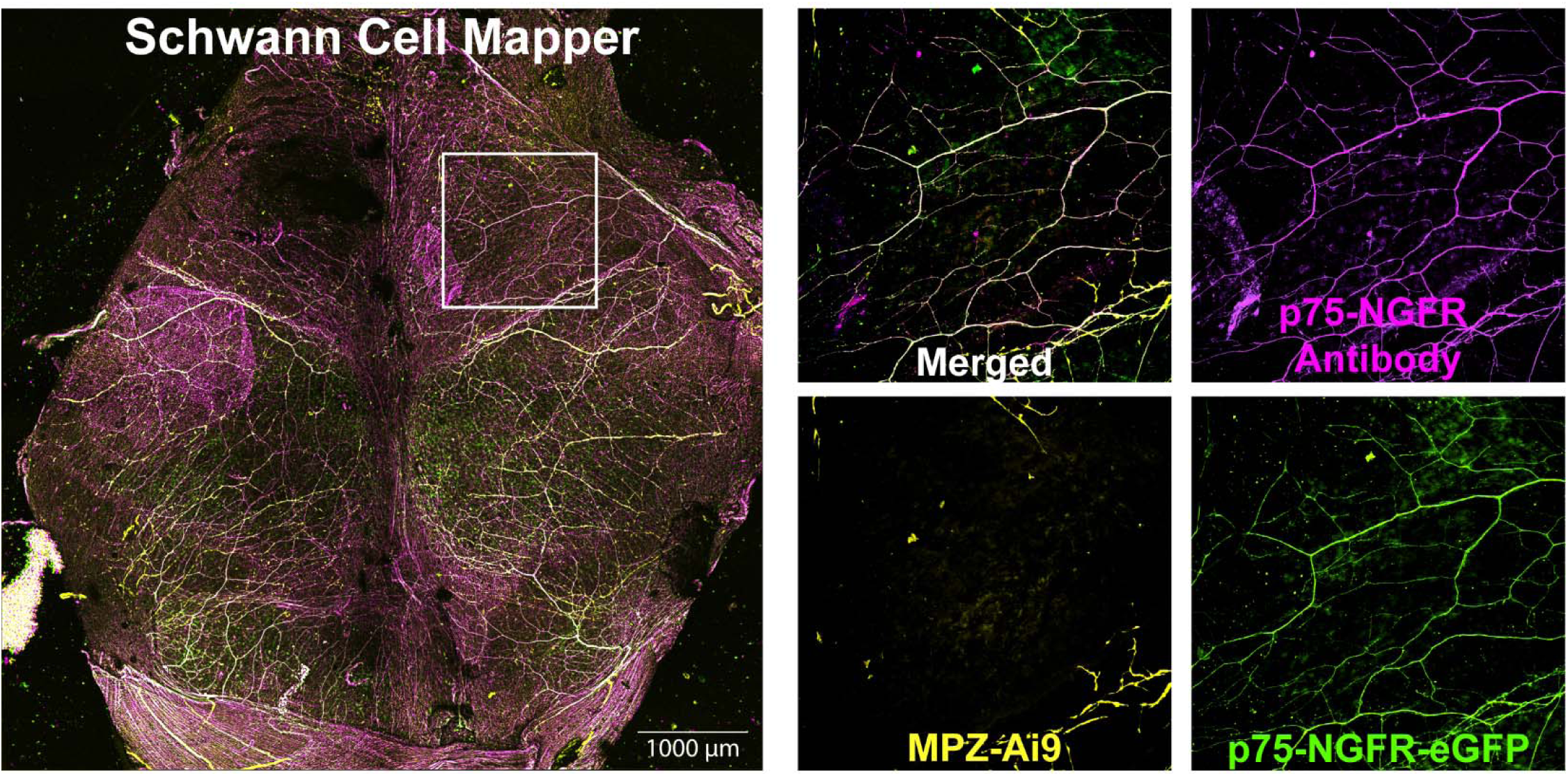
p75-NGFR+ and MPZ+ cells in the mouse calvaria at birth. Maximum intensity projection of 4x spinning disk confocal images of ‘Schwann Cell Mapper’ (*Mpz*-Cre^+/-^;TdT^+/+^;*Ngfr*-eGFP mouse calvaria showing p75-NGFR+ and MPZ+ SCs, and colocalization with p75-NGFR immunolabeling at Day 0 post birth. Scale = 1000 µm. The white box indicates a zoomed in portion to the right. Magenta = p75-NGFR antibody, Yellow = MPZ-Ai9 reporter for myelinating SCs, Green = p75-eGFP reporter for non-myelinating SCs.

### MPZ and p75-NGFR label two distinct populations of Schwann cells

MPZ and p75-NGFR are thought to label two distinct SC populations. To assess this in bone, we imaged the calvarial periosteum of adult ‘Schwann Cell Mapper’ mice at 40x resolution (Fig.5A). This showed that nerve bundles within the periosteum were often made up of multiple small axons and SCs running together. Within these bundles, MPZ and p75-NGFR generally labeled two different populations of SCs. However, in rare cases, nerve-lining SCs expressed both MPZ and p75-NGFR simultaneously (Fig.5A). This separation between MPZ and p75-NGFR was difficult to appreciate at lower resolutions given the small size and close relationship between SCs and axons within periosteal nerves, making co-expression appear more prevalent than occurred at high resolution.

**Figure 5.**
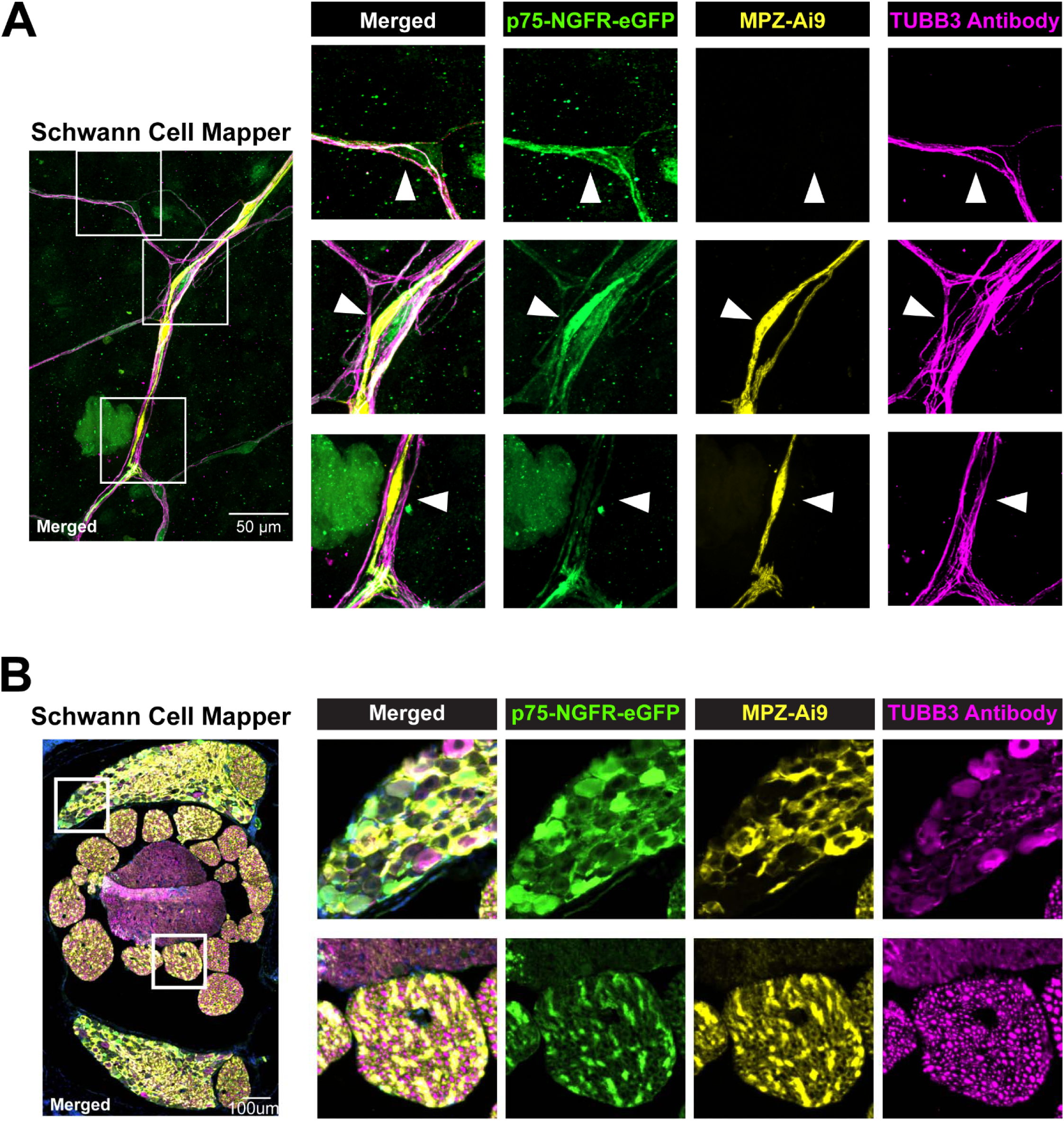
Co-localization and specificity of p75-NGFR and MPZ reporters. **(A)** Maximum intensity projection (MIP) of whole mount 40x spinning disk confocal images of a ‘Schwann Cell Mapper’ (*Mpz*-Cre^+/-^;TdT^+/+^;*Ngfr*-eGFP^+/+^) mouse calvaria showing coexpression of both p75-NGFR-eGFP and MPZ-Ai9 reporters within a Schwann cell body while maintaining association with TUBB3 antibody, a nerve marker. White boxes indicate zoomed in portions to the right. Scale = 50 µm. **(B)** MIP of 20x spinning disk confocal images of a ‘Schwann cell mapper” mouse spine sections showing expression of p75-NGFR-eGFP and MPZ-Ai9 separate from TUBB3+ axons in the Dorsal Root Ganglia. White boxes indicate zoomed in portions to the right. Scale = 50 µm. Magenta = TUBB3 antibody, Green = p75-NGFR-eGFP reporter for non-myelinating SCs, Yellow = MP0-Ai9 reporter for myelinating SCs.

Nerves may also express p75-NGFR cell autonomously, though this is thought to be limited to developing axons.(Barrett and Bartlett, 1994) To check this, we analyzed cross sections of the spinal cord from ‘Schwann Cell Mapper’ mice containing neural cell bodies in the dorsal root ganglia (DRG) and descending axon tracts. Some cell bodies in the DRG were positive for p75-NGFR. However, axon tracts generally showed punctate TUBB3^+^ axon staining that had limited overlap with MPZ and p75-NGFR, implying minimal axon-level expression of these reporters in healthy adult animals (Fig.5B). Instead, MPZ lineage^+^ and p75-NGFR^+^ cells either wrapped the TUBB3^+^ axons, consistent with SCs, or were located between clusters of nerve fibers.

### MPZ and p75-NGFR Schwann cells are most prevalent in periosteum and enter bone marrow through cortical canals

In skeletal tissues, the periosteum contains the highest density of nerve axons.(Lorenz et al., 2021; Steverink et al., 2021) Nerves then enter bone alongside blood vessels through transcortical canals and distribute within the bone marrow.(Chen et al., 2007; Lorenz et al., 2021) Consistent with this, MPZ lineage^+^ and p75-NGFR^+^ SCs were most prevalent within the periosteum as visualized in the calvaria, caudal vertebrae, femur, and tibia (Fig.6A-F, Supplemental Fig.2). In the mouse calvaria, the sutures were also highly innervated and contained abundant SCs (Fig.6A,B). Like nerves, SCs also entered bone through transcortical canals (Fig.6A-F). In some cases, nerve-lining SCs could be observed in close approximation to trabecular bone, though this was relatively uncommon (Fig.6E, Supplemental Fig.2).

**Figure 6.**
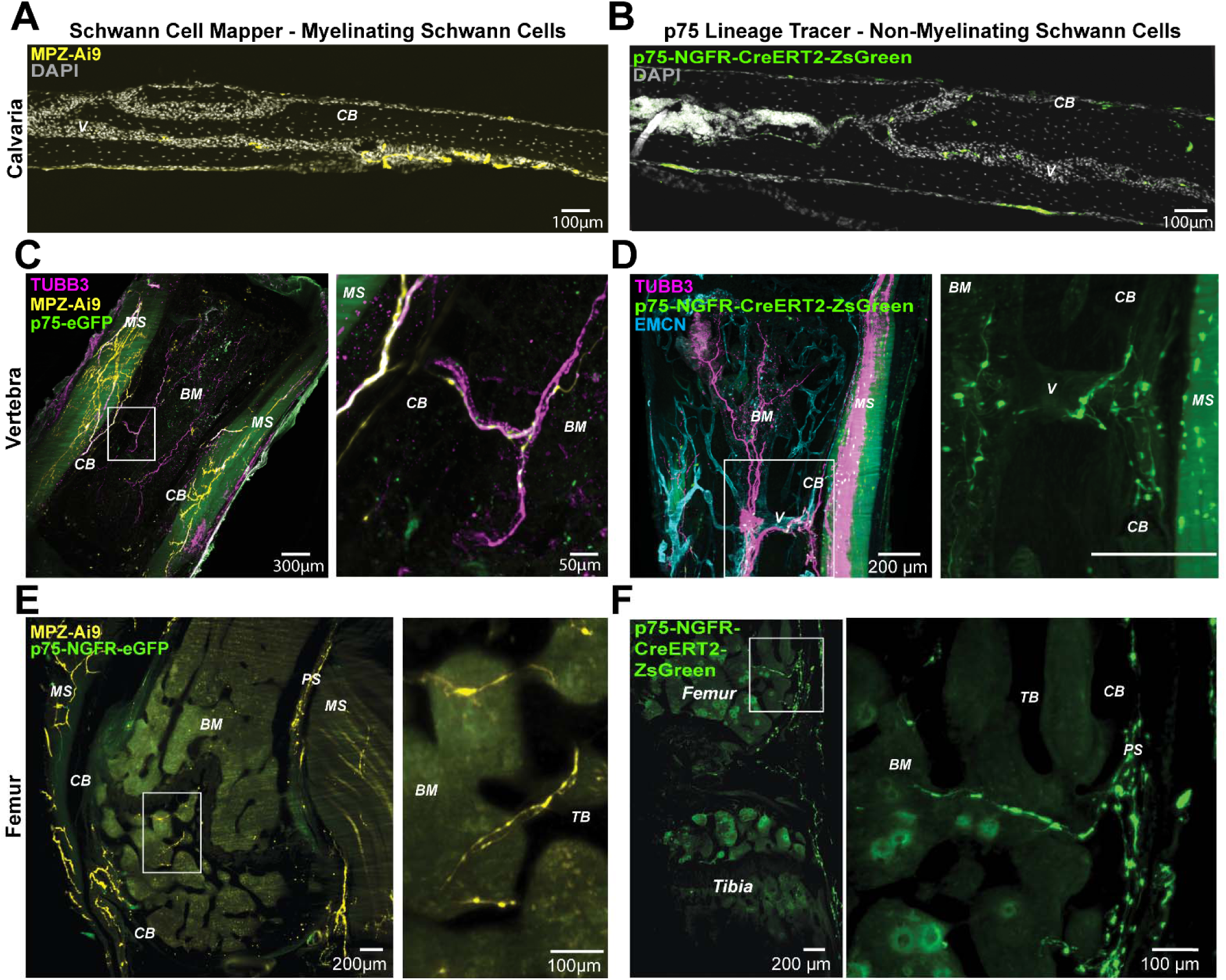
Schwann cells enter the bone marrow space from the periosteum via cortical canals. **(A)** Calvaria section of a ‘Schwann Cell Mapper’ (*Mpz*-Cre^+/-^;TdT^+/+^;*Ngfr*-eGFP^+/+^) mouse showing MPZ+ Schwann cells (SCs) along a blood vessel. Scale = 100 µm **(B)** Calvaria section of a ‘p75 Lineage Tracer’ (*Ngfr*-CreERT2^+/-^;ZsGreen1^+/+^) mouse showing p75-NGFR-eGFP+ cells along a blood vessel. Scale = 100 µm **(C)** Maximum intensity projection (MIP) of 4x lightsheet microscopy images of ‘Schwann cell mapper’ mouse caudal vertebra showing the abundance of myelinating MPZ+ SCs in the muscles and periosteum. In addition, MPZ+ SCs enter the vertebra bone marrow space along with TUBB3+ nerves. Scale = 300 µm. The white box indicates a zoomed in portion to the right. (**D)** MIP of lightsheet microscopy images of ‘p75 Lineage Tracer’ mouse caudal vertebra showing non-myelinating p75-NGFR+ SCs entering the bone marrow space along with EMCN+ blood vessels and TUBB3+ nerves. The white box indicates a zoomed in portion to the right. Scale = 200 µm. **(E)** MIP of lightsheet images of a ‘Schwann cell Mapper’ mouse femur showing the greater proportion of myelinating MPZ+ SCs present in the periosteum and muscle surrounding the bone compared to the bone marrow space. Scale = 200 µm. The white box indicates a zoomed in portion to the right and shows myelinating SCs entering the subchondral bone compartment. Scale = 100 µm. **(F)** MIP of lightsheet images of ‘p75 Lineage Tracer’ mouse knee joint showing distribution of non-myelinating SCs in the femur and tibia periosteum and the knee fat pad. Scale = 200 µm. The white box indicates a zoomed in portion to the right and shows that non-myelinating SCs can also enter the bone marrow space of long bone from the periosteum. Scale= 100µm. Gray = DAPI, Yellow = MPZ-Ai9 reporter for myelinating SCs, Green = p75-eGFP or p75-NGFR-CreERT2-ZsGreen as indicated, Blue = EMCN. CB= Cortical Bone, TB= Trabecular Bone, BM= Bone Marrow, MS= Muscle, PS=periosteum, V = Blood Vessel.

### MPZ and p75-NGFR Schwann cells have unique branching patterns

In cats, a single axon can innervate a 2 to 4 mm^2^ region of periosteum.(Mahns et al., 2006) This implies extensive branching of individual axons that would require a large population of SCs with high network forming capacity. To define the size and morphology of SCs in mouse bone, we reviewed individual SCs within calvaria imaging datasets. For this analysis, we focused on the MPZ-Cre;Ai9 reporter in the ‘Schwann Cell Mapper’ and the p75-NGFR-CreERT2;ZsGreen reporter in the ‘p75 Lineage Tracer’ as these fluorophores were very bright in both the cytoplasm and the nucleus, making it easy to identify individual cells.

Within the calvarial periosteum, most MPZ lineage SCs had limited branching, extending predominantly along one nerve fiber (Fig.7A). Linear MPZ lineage SCs presented as either single cells or grouped into cords (Fig.7A), reflecting the presence of multiple axons running in parallel. When grouped, the nuclei of the individual MPZ lineage SCs were aligned and offset by a few µm each in a diagonal pattern (Fig.7A). In some cases, MPZ lineage SCs had accessory branches or sprouts that extended from the proximal or distal cytoplasm, or as a third projection near to the nucleus (Fig.7A). In the tibia, MPZ lineage SCs with a linear pattern were also observed primarily in the periosteum (Fig.7B). By contrast, branching was prevalent in the tibial bone marrow with formation of intricately branched structures in some regions of the diaphysis (Fig.7B).

**Figure 7.**
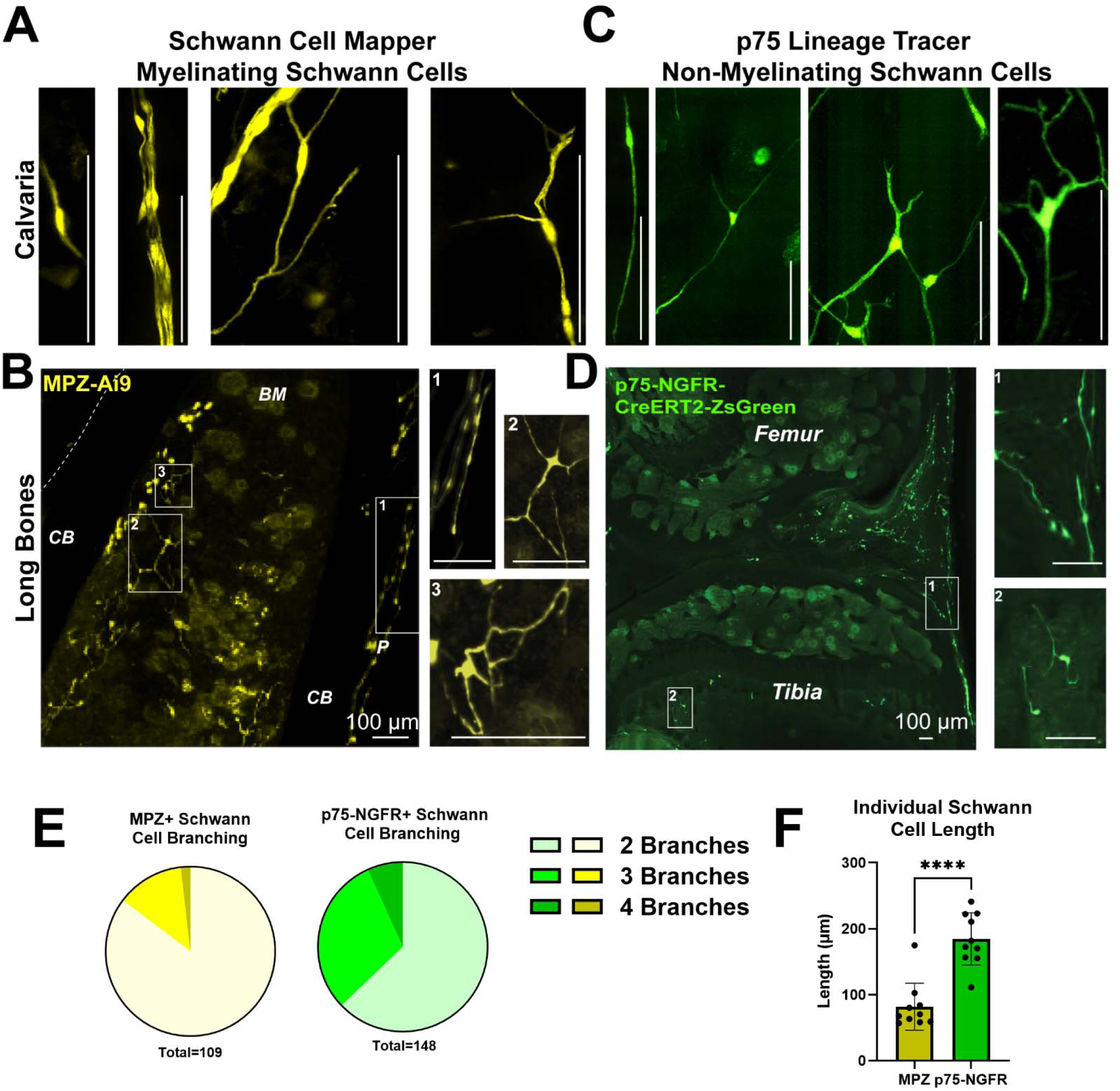
Morphologies of myelinating and non-myelinating Schwann cells. Selections from **(A)** whole mount spinning disk confocal images and **(B)** lightsheet images of ‘Schwann Cell Mapper’ (*Mpz*-Cre^+/-^;TdT^+/+^;*Ngfr*-eGFP^+/+^) mouse calvaria and long bones representative of the breadth of morphologies of myelinating MPZ+ SCs in the periosteum and marrow space. Scale = 100µm Selections from **(C)** whole mount spinning disk confocal images and **(D)** lightsheet images of ‘p75 Lineage Tracer’ (*Ngfr*-CreERT2^+/-^;ZsGreen1^+/+^) mouse calvaria and long bones representative of the breadth of morphologies of non-myelinating SCs. Scale = 100µm. **(E)** Quantification of branching of MPZ+ and p75-NGFR+ SCs in mouse calvariae (total N represents individual SCs). **(F)** Quantification of the length of individual MPZ+ and p75-NGFR+ SCs from the frontal and parietal bones (total length of the SC filaments divided by the number of nuclei along the SC filaments, dots represent individual mice). Yellow = MPZ-Ai9 reporter for myelinating SCs. Green = p75-NGFR-CreERT2-ZsGreen reporter for non-myelinating SCs. CB = Cortical Bone, BM = Bone Marrow, P = periosteum.

In the calvaria, p75-NGFR^+^ SCs commonly presented with 3 to 4 individual branches that extended away from the nucleus as a central node (Fig.7C). These branches could split into smaller offshoots to develop additional complexity (Fig.7C). Similar branching of p75-NGFR+ SCs was also observed in the long bones (Fig.7D). When quantified within the calvarial periosteum, the proportion of SCs with 3+ branches was 15% for MPZ lineage^+^ cells and 45% for p75-NGFR^+^ cells (Fig.7E).

Similarly, the estimated length of p75-NGFR^+^ SCs in the calvarial periosteum was 2.26x greater than MPZ lineage^+^ SCs (184.3+37.6 µm vs 81.7+33.8 µm, p<0.0001) (Fig.7F). Overall, non-myelinating SCs in periosteum were longer with more extensive branching than myelinating SCs.

### MPZ and p75-NGFR also label fibroblast-like cells in periosteum and bone marrow with features of immature Schwann cells

In addition to labeling mature SCs, MPZ and p75-NGFR can also be expressed by SCPs and iSCs (NCSC ➔ SCP ➔ iSC ➔ mature SC). In calvarial periosteum of adult mice, we identified two distinct populations of large, flat cells with a dense central nucleus and thin spreading cytoplasm (“fried egg” type morphology) that were either MPZ lineage^+^ or p75-NGFR^+^ (Fig.8A-D). In general, individual flat cells were distributed throughout the calvarial periosteum in between nerve fibers and mature SCs. The flat cells differed from traditional SCs as they were not directly associated with TUBB3^+^ nerves and lacked “bead on a string” morphology. In some cases, it appeared that transitioning flat cells were starting to become associated with TUBB3^+^ nerves, consistent with maturation of iSCs to mature SCs (Supplemental Figure 3).

**Figure 8.**
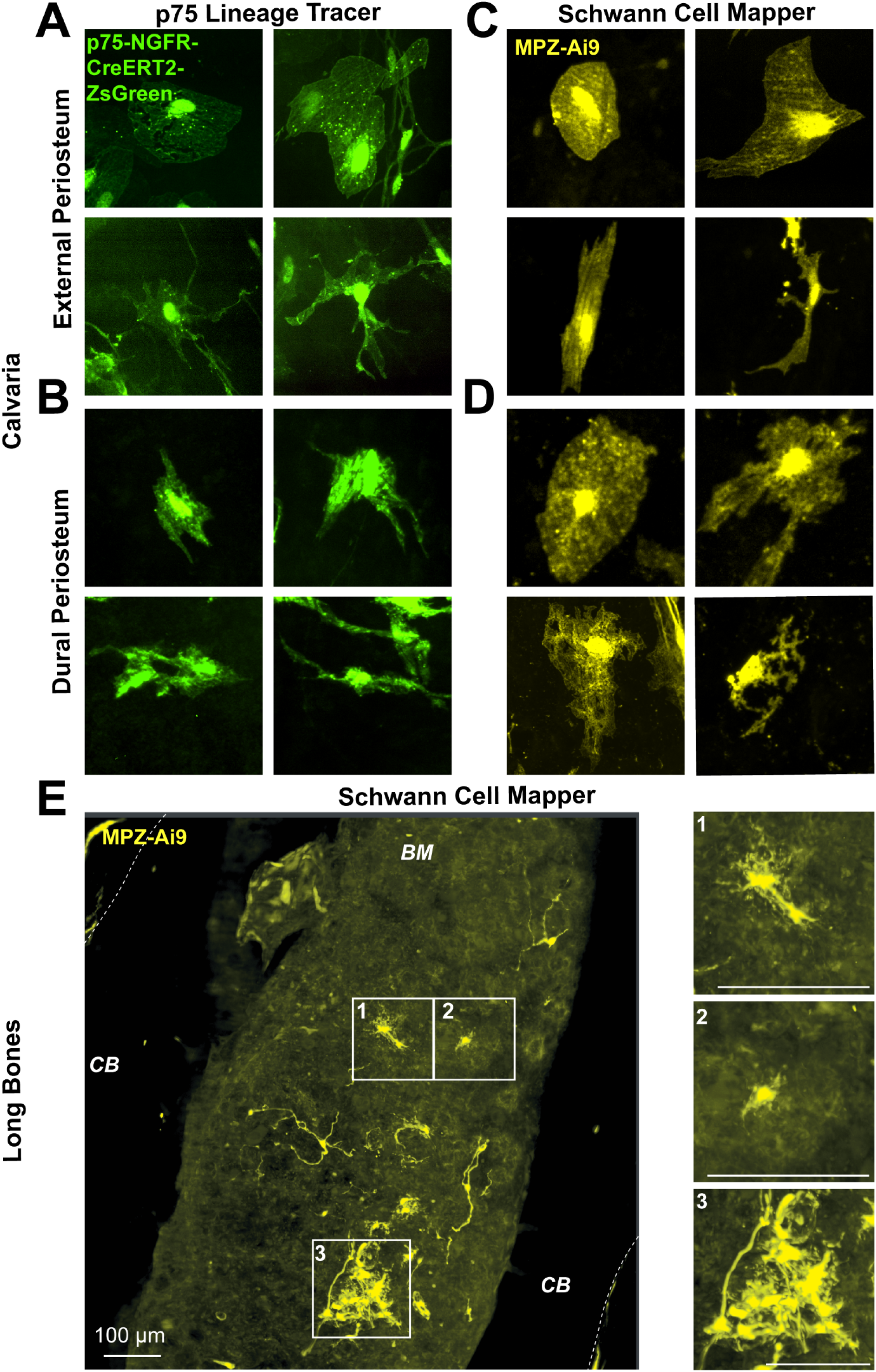
Candidate immature Schwann cell morphologies of MPZ+ and p75+ cells. **(A)** Selections from whole mount spinning disk confocal images of ‘p75 Lineage Tracer’ (*Ngfr*-CreERT2^+/-^;ZsGreen1^+/+^) mouse calvariae representative of the breadth of morphologies of p75+ candidate immature Schwann cells (iSCs) in the external periosteum and **(B)** in the dural periosteum. **(C)** Selections from whole mount spinning disk confocal images of ‘Schwann Cell Mapper’ (*Mpz*-Cre^+/-^;TdT^+/+^;*Ngfr*-eGFP^+/+^) mouse calvariae representative of the breadth of morphologies of MPZ+ iSCs in the external periosteum and **(D)** in the dural periosteum. **(E)** Selections lightsheet microscopy images of ‘Schwann Cell Mapper’ mouse long bone representative of the breadth of morphologies of MPZ+ cells in the bone marrow compartment. Scale = 100µm. Yellow = MPZ-Ai9 reporter for myelinating SCs. Green = p75-NGFR-CreERT2-ZsGreen reporter for non-myelinating SCs. CB= Cortical Bone, BM= Bone Marrow.

Within the calvarial periosteum, the MPZ lineage^+^ and p75-NGFR^+^ flat cells ranged from round with a high cytoplasm to nucleus ratio, to oblong and elongated with maximum diameters of 20 to 80 µm (Fig.8A-D). Some also had fibroblast or SC like projections with lengths from 10 to 40 µm (Fig.8A-D). P75-NGFR^+^ flat cells on the dural periosteum had a more uniform spindle-like morphology than on the external periosteal surface (Fig.8C). Though most presented as individual cells, occasional clusters containing 3 to 30 cells were observed. MPZ lineage^+^ and p75-NGFR^+^ flat cells were present in areas with nerves both close to and far from the suture. Similar MPZ lineage^+^ cells with iSC morphology and occasional cell clusters were found in the bone marrow of the tibia (Fig.8E). We were not able to identify MPZ lineage^+^ or p75-NGFR^+^ candidate iSCs in the periosteum of the light sheet datasets (femur, tibia, vertebrae) but suspect that this may be due to limitations of the imaging rather than true absence.

### Quantification of MPZ and p75-NGFR mature SCs and candidate iSCs reveals distinct periosteal niches in neural crest vs mesoderm-derived calvarial bone

Next, we quantified the total length and number of periosteal MPZ lineage^+^ and p75-NGFR^+^ mature SCs and candidate iSC-like flat cells, respectively. To consider possible effects of developmental origin, quantifications were compared between neural crest-derived frontal bone and mesoderm-derived parietal bones of male and female mice at 12 to 24 weeks of age (Fig.9A,B).

**Figure 9.**
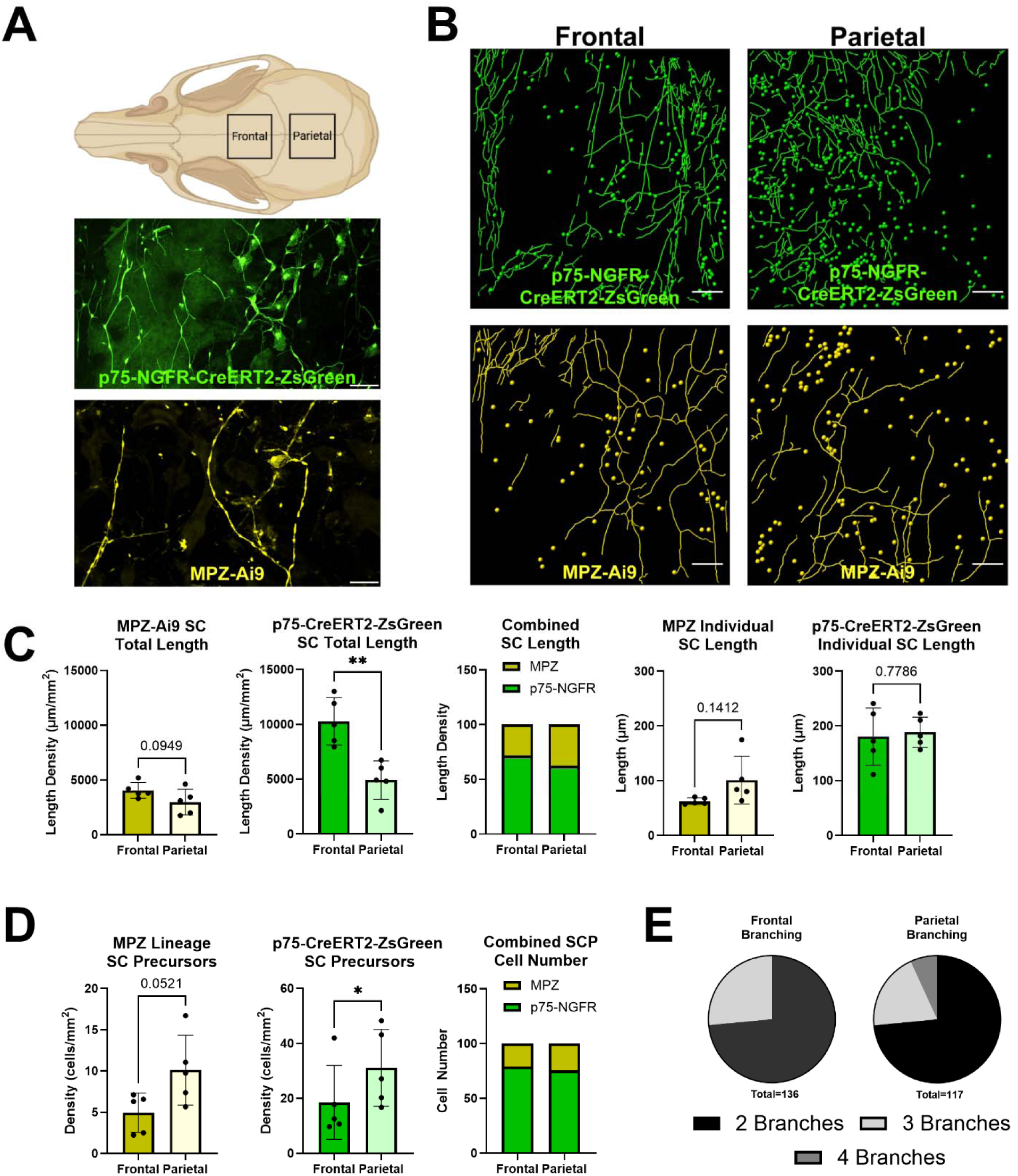
Differences in the periosteal niche of the mouse calvaria between frontal and parietal bones. **(A)** Maximum intensity projection of whole mount 10x spinning disk confocal images of a ‘Schwann Cell Mapper’ (*Mpz*-Cre^+/-^;TdT^+/+^;*Ngfr*-eGFP^+/+^) mouse calvaria and a ‘p75 Lineage Tracer’ (*Ngfr*-CreERT2^+/-^;ZsGreen1^+/+^) mouse calvaria representative of the patterns and distribution of mature Schwann cells (SCs) and candidate immature SCs. Yellow = MPZ-Ai9 reporter for myelinating SCs. Green= p75-NGFR-CreERT2-ZsGreen reporter for non-myelinating SCs. **(B)** Representative Masks of frontal and parietal bones of the ‘p75 Lineage Tracer’ and ‘Schwann Cell Mapper Mouse. Lines = mature SCs, Dots = candidate iSCs. Scale = 200 µm. **(C)** Quantification of MPZ+ and p75-NGFR+ SCs total length and individual length in frontal and parietal bones (dots on graphs represent individual mice). **(D)** Quantification of MPZ+ and p75-NGFR+ candidate iSC number and ratio in frontal and parietal bones (dots on graphs represent individual mice). **(E)** Quantification of branching of SCs in frontal and parietal bones (total N represents individual SCs).

Neural crest-derived frontal bone had 1.93x more p75-NGFR^+^ mature SC length and 1.35x more, but not statistically significant, MPZ lineage^+^ mature SC length than the parietal bone (Fig.9C). By contrast, there were 2.06x more MPZ lineage^+^ and 1.68x more p75-NGFR^+^ candidate iSCs in the parietal bone than frontal bone (Fig.9D). Despite these differences in SC maturity, the ratio of MPZ to p75-NGFR SC length and candidate iSC number was relatively consistent between the frontal and parietal bones (Fig.9E). The length of individual mature SCs (Fig.9F) and the percentage of mature SCs with 3 or 4 branches was also comparable between sites (Fig.9G). Overall, within calvarial periosteum, there was more non-myelinating SC length and p75-NGFR^+^ iSC number at a ratio of ∼2:1 and ∼3:1 relative to MPZ lineage^+^ populations, respectively.

### MPZ-Cre labels rare osteocytes, osteoblasts, adipocytes, and chondrocytes in healthy adult mice

In the ‘Schwann Cell Mapper’ model, MPZ-Cre is constitutively expressed, meaning that any cell derived from an MPZ-expressing cell during development will be positive for Ai9, an ultrabright TdTomato reporter. The Ai9 reporter is not dose dependent, it is either on or off. For example, if all osteoblasts were derived from an MPZ^+^ progenitor lineage, we would expect to see bright expression of Ai9/TdTomato in adult osteolineage cells that matches that of MPZ lineage SCs. We reviewed neural crest-derived frontal and interparietal bones and mesoderm-derived parietal, femur, tibia, and vertebrae bones for evidence of MPZ lineage cells. Bone-lining cells (osteoblasts/clasts) and osteocytes were generally negative for MPZ-Cre;Ai9 in all bones examined, regardless of developmental origin. However, in rare cases MPZ lineage^+^ osteoblasts and osteocytes that co-expressed osterix (OSX) were detected (Fig.10A-B). In a review of 1,000 osteocytes across 3 adult mice at 12 to 14 weeks of age, only 3/1000 cells were positive for MPZ-Ai9 (0.03%). Overall, we estimate that labeling occurred in <0.1% of osteolineage cells in all bones examined across cortical and trabecular compartments with no apparent pattern or relationship to the developmental origin of the bone examined (Fig.10C). Rare MPZ lineage^+^ bone marrow adipocytes and chondrocytes were also detected in long bone and vertebrae, suggesting potential for trilineage differentiation of MPZ^+^ cells. However, though possible, this does not appear to be a major contributor to osteoblast, adipocyte, and chondrocyte lineages in healthy adult mice.

### P75-NGFR reporter expression is largely limited to SCs and iSCs, with rare expression in other bone cell types

P75-NGFR expression has been reported in skeletal cell lineages including pericytes, mesenchymal stem cells, osteoblasts, and even osteoclasts.(Huber and Chao, 1995; Mikami et al., 2012; Chartier et al., 2017; Luo et al., 2025) However, this has been based on gene expression studies or antibodies, which are subject to technical limitations and overinterpretation, making the *in vivo* reality on a per cell basis unclear. In our reporter models, there was limited evidence of p75-NGFR expression in bone outside of SCs and candidate iSCs in healthy adult mice. In the ‘p75 Lineage Tracer’ model, tamoxifen is provided to activate Cre-dependent expression of ZsGreen 48-hours prior to analysis (on/off system). This means that adult cells actively expressing p75-NGFR will have bright intensity of ZsGreen. In the ‘Schwann Tracer’ model, p75-eGFP also marks cells actively expressing p75-NGFR. As for MPZ, osteocytes and osteoblasts were generally negative for p75-NGFR (Fig.1,3,4,6,7,9), apart from a few scattered osteocytes and osteoblasts that co-expressed MPZ and P75-NGFR (Fig.10, estimated as <0.1% of cells). No p75-NGFR+ bone marrow cells, osteoclasts, adipocytes, or chondrocytes were observed. However, in some samples a rare population of p75-NGFR^+^ square, flat cells were found to wrap vascular-appearing structures in calvarial bone (Supplemental Fig.4). These were not common or present in all mice. When observed, the flattened morphology of these cells was different than traditional spindle-shaped pericytes, and tended to resemble the large, flat candidate iSCs with high cytoplasm:nucleus ratio discussed above.

**Figure 10.**
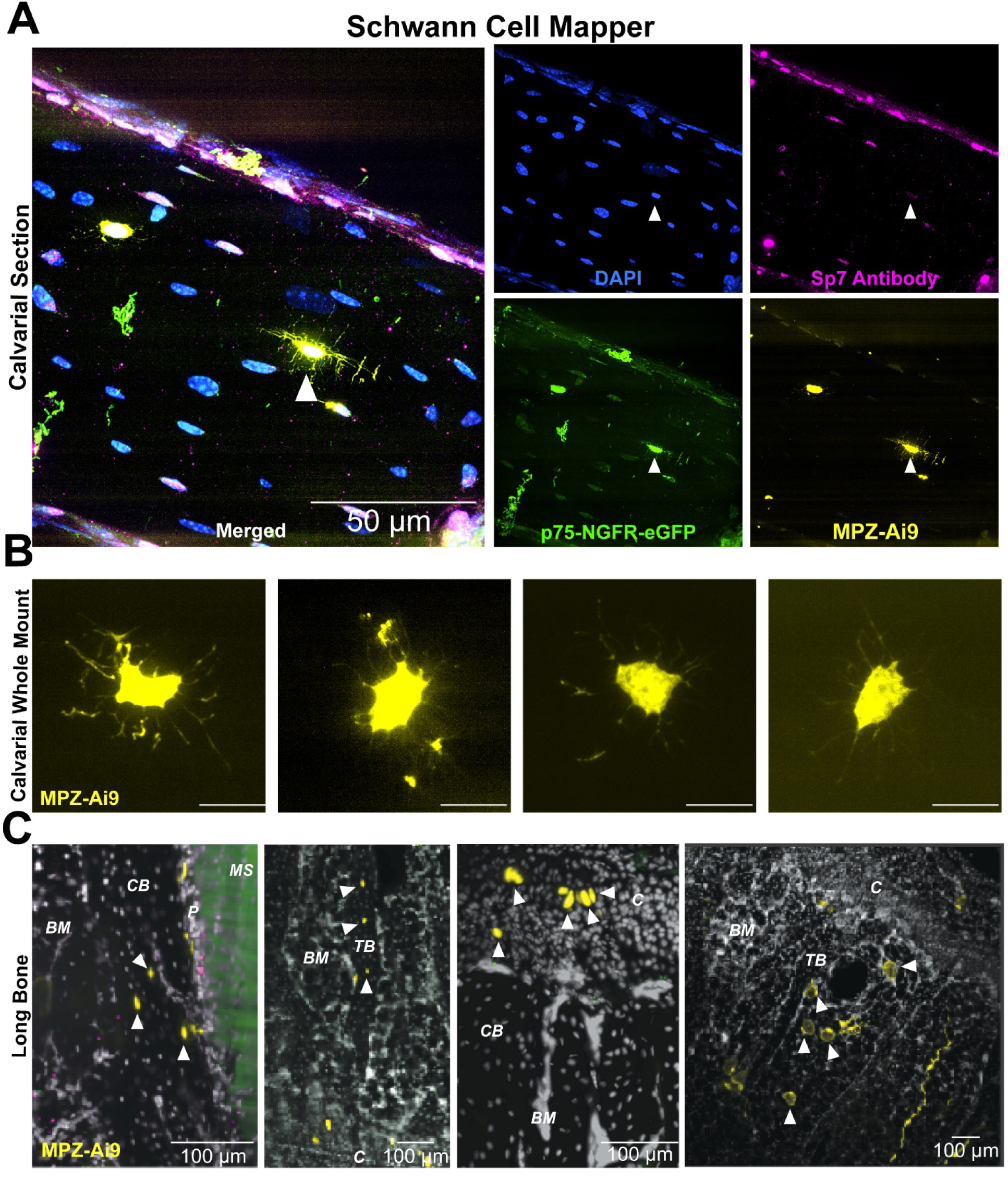
Evidence of rare p75-NGFR+ and MPZ+ osteolineage, cartilage, and bone marrow adipocyte cells. **(A)** 100x spinning disk confocal images of ‘Schwann Cell Mapper’ (*Mpz*-Cre^+/-^;TdT^+/+^;*Ngfr*-eGFP^+/+^) mouse calvaria sections showing p75-NGFR+ and MPZ+ osteocytes. Scale=50 µm. Magenta= SP7 antibody. Yellow= MPZ-Ai9 reporter. Green = p75-NGFR-eGFP reporter. Blue= DAPI. **(B)** Selections of whole mount 60x spinning disk confocal images of ‘Schwann Cell Mapper’ mouse calvariae showing representative MPZ+ osteocytes. Scale = 15 µm. Yellow = MPZ-Ai9 reporter. **(C)** Selections of lightsheet microscopy images of ‘Schwann Cell Mapper’ mouse caudal vertebrae showing rare osteolineage cells within the cortical bone, trabecular bone, bone marrow, and cartilage. White Arrows indicates MPZ+ osteolineage, cartilage, or bone marrow adipocyte cells. Scale = 100 µm. Yellow = MPZ-Ai9 reporter. Green = p75-NGFR-eGFP reporter. Gray= DAPI. CB= Cortical Bone, TB= Trabecular Bone, BM= Bone Marrow, MS= Muscle, C= Cartilage.

### Survey of MPZ and p75-NGFR in other tissues

In a final imaging study, we also completed a survey of several peripheral tissues to inform future investigators that wish to utilize the ‘Schwann Cell Mapper’ and ‘p75 Lineage Tracer’ models. This included the heart, liver, intestine, kidney, pancreas, and spleen (Supplemental Fig.5). In the intestine, pancreas, and spleen, there was classical SC morphology which was highly associated with TUBB3^+^ nerves. However, we also observed MPZ lineage^+^ and/or p75-NGFR^+^ cells with morphology differing from SCs. In the heart, single MPZ lineage^+^ cells were interspersed throughout the cardiac muscle. The morphology, size (10-100 µm long x 5-20 µm wide), and distribution was consistent with labeling a subset of cardiomyocytes. Very small, nucleated p75-NGFR^+^ cells were also occasionally present in between the cardiac muscle fibers. In the liver, there were individual MPZ lineage^+^ and/or p75-NGFR^+^ hepatocytes occasionally present. There was also a robust distribution of p75-NGFR^+^ cells between the hepatocytes with morphology consistent with prior reports of p75-NGFR^+^ hepatic stellate cells.(Trim et al., 2000). In the intestine, individual MPZ lineage^+^ cells, p75-NGFR^+^ cells, or cells expressing both reporters were interspersed throughout the intestinal epithelium. In the kidney, there was a rare population of small, variably shaped MPZ lineage^+^ or p75-NGFR^+^ cells scattered through the tissue. In the pancreas, there were many p75-NGFR^+^ SCs associated with TUBB3^+^ nerves, especially entering the duct. We also observed a few small, round MPZ lineage^+^, p75-NGFR^+^ cells throughout the pancreas that were not associated with TUBB3^+^ nerves. Lastly, the spleen had a large population of p75-NGFR^+^ fibroblast-like cells in clusters with small, round MPZ lineage^+^ cells interspersed.

## DISCUSSION

Herein, we present an imaging-based framework of SC subtypes in bones of different embryonic origins. Previous studies have shown that MPZ and p75-NGFR are unique markers of myelinating and non-myelinating SCs, respectively, in nerves like the sciatic nerve(Yasuda et al., 1987; Bentley and Lee, 2000), and we confirm this labelling strategy in adult mouse bone, bone marrow, and periosteum. This effectively marks SCs in the calvaria, both mesoderm and neural crest derived bone, as well as SCs in other mesoderm derived bones, including the tibia, femur, and vertebrae. For the first time, we also identify candidate populations of both MPZ lineage^+^ and p75-NGFR^+^ iSCs throughout the calvarial periosteum and find that bones of different embryonic origins have altered abundance of candidate iSCs relative to mature SCs. Additional observations are detailed below.

### Nerves in bone are covered by mature myelinating and non-myelinating SCs, creating an adaptable interface between nerve and bone

This study shows that both MPZ lineage^+^ myelinating and p75-NGFR^+^ non-myelinating SCs cover nerves throughout the periosteum, bone, and bone marrow and enter the bone through transcortical canals. Consistent with this, the relative density of SCs mirrored that of peripheral nerves(Brazill et al., 2019; Lorenz et al., 2021), being highest in periosteum and lower in bone and bone marrow. Classically, mature SCs are thought to either be myelinating or non-myelinating, expressing separate markers like MPZ and p75-NGFR respectively.(Yasuda et al., 1987; Bentley and Lee, 2000) Most of the cells we observed in bone held this pattern, although surprisingly, we found some mature, nerve-lining SCs that co-expressed both MPZ and p75-NGFR. This population was rare and was not associated with any specific location. This dual labeled population may represent SCs in transition, from non-myelinating to myelinating or vice versa, due to changes in associated axon size or signaling. This potential plasticity of mature SCs, most often through a repair SC intermediate, has been documented previously but remains poorly understood *in vivo.*(Gomez Sanchez et al., 2017)

Non-myelinating SCs in calvarial periosteum were longer than myelinating SCs (184 µm vs 82 µm on average). Non-myelinating SCs also had evidence of increased branching vs myelinating SCs which tended to only have two branches, extending their processes a short distance along an axon, with many SCs in succession along the axon length. This is somewhat different than previous literature in larger nerve trunks such as sciatic nerve which shows myelinating SCs to be much longer than non-myelinating SCs, 500-800 µm vs 100-350 µm (Gomez Sanchez et al., 2017) and suggests unique adaptations of SCs within terminal end organs such as bone.

Overall, these findings reinforce the concept that mature SCs represent an adaptable cellular interface between nerve axons and nearby skeletal cells. Mature SCs are surrounded by a basal lamina,(Webster, 1971) a specialized layer of extracellular matrix that can act as a semipermeable barrier, structural scaffold, and anchoring site between tissues. SCs also interact directly with their encircled axon(s) through gap junctions, paranodal junctions, ion channels, and secretion of extracellular vesicles.(Samara et al., 2013) In addition, SCs can both produce and respond to secreted signaling factors.(Samara et al., 2013; Oliveira et al., 2023) Altogether, SCs are structurally positioned along nerves in bone with unique adaptations that may modulate nerve sprouting and nerve function in skeletal tissues. SCs also have high potential to regulate niche-level interactions between local nerves and bone cells, influencing aspects of bone pain, homeostasis, and healing.(Wei et al., 2019; Ikami et al., 2020; Zhang et al., 2023)

### Periosteum contains a robust population of candidate iSCs

Nerve sprouting during development drives the maturation of SCPs to iSCs to mature SCs through signals including NRG1.(Leimeroth et al., 2002; Taveggia et al., 2005) The primary distinction between SCPs and iSCs is that iSCs produce their own basal lamina and do not depend on axonal contact or axon-derived cues for survival, but rather acquire autocrine survival loops.(Webster, 1971; Bunge and Bunge, 1983; Meier et al., 1999) There is evidence that SCPs can persist in adult tissues including mouse incisors and guts of fish.(Kaukua et al., 2014; El Nachef and Bronner, 2020) However, the presence of iSCs and SCPs in other adult organs remains an active and controversial area of research.

In this study, we observed two distinct populations of large, MPZ lineage^+^ and p75-NGFR^+^ flat cells interspersed within richly innervated areas of periosteum. Cell morphology was “fried egg-like” with a dense central nucleus and thin spreading cytoplasm, ranging from rounded with high cytoplasm to nucleus ratio to somewhat more spindle shaped with fibroblast-or SC-like projections. There was also evidence of transitional appearing cells (between flat iSC and mature SC) undergoing nerve axon association. Overall, the reporter expression, cell morphology, localization relative to nerve axons, and lack of direct axon contact is consistent with a periosteum-resident population of iSCs rather than axon-dependent SCPs. Some evidence for the presence of a limited candidate population of iSCs in bone marrow was also observed, particularly with the MPZ lineage tracer.

However, more work will be needed to clarify this in future publications.

The primary implications of a resident iSC population in bone are twofold. First, a network of periosteal iSCs may support local neurogenesis and axon plasticity. Though there are exceptions, SCs and iSCs classically coordinate axon sprouting, extending processes ahead of new nerves to create a cellular track.(Hromada et al., 2024) Both SCs and iSCs can also secrete axon guidance molecules that act as chemoattractants for new sprouts.(Fontana et al., 2012; Brushart et al., 2013) Second, iSCs may support bone homeostasis and healing through paracrine signals and possibly also undergo direct differentiation to mesenchymal lineages.(Kaukua et al., 2014) Based on current literature and our lineage tracing results, trophic support by SC lineages seems more prevalent than direct differentiation.(Zhang et al., 2021; Yang et al., 2026) However, this is a new and emerging area of research, and additional work will be needed to clearly delineate the contributions of iSCs, mature SCs, and repair SCs to bone health.

### Neural crest-derived frontal bone favors mature SCs while mesoderm-derived parietal bone favors candidate iSCs

We characterized the differences in candidate iSC number and mature SC length between the frontal and parietal bones, which derive from neural crest and mesoderm embryonic origins, respectively. Although present in both, mesoderm-derived parietal bone contained more p75-NGFR^+^ and MPZ lineage^+^ flat cells (candidate iSCs) than neural crest-derived frontal bone. Conversely, mature SCs were enriched in neural-crest derived frontal bone. This could be due to altered patterning during development, or differences within the periosteal signaling niche of adult mice that remain to be defined. Similar to mature SCs, there were two separate subpopulations of MPZ lineage^+^ and p75-NGFR^+^ candidate iSCs, with very few dual labeled cells. It is possible that cells labeled with MPZ but not p75-NGFR may be further along the differentiation pathway. It is also possible that the MPZ lineage^+^ and p75-NGFR^+^ iSC-like c are pre-primed to form mature myelinating and non-myelinating SCs, respectively. Overall, iSCs are poorly characterized in peripheral tissues and this remains an open point of future investigation in bone and beyond.

### MPZ and p75-NGFR lineage cells show trilineage potential

NCSCs, SCPs, and potentially also iSCs and repair SCs have been proposed to give rise to multiple cell lineages, particularly *in vitro*.(Adameyko et al., 2009; Furlan et al., 2017; Lumb et al., 2018; Xie et al., 2019) Our *in vivo* lineage tracing data are consistent with this potential in that we observed MPZ lineage^+^ osteocytes, adipocytes, and chondrocytes in the adult skeleton. However, these events were infrequent (<0.1% of cells) and may reflect lineage contributions established during development rather than ongoing plasticity in adulthood. Previous literature using an MPZ-Cre model reported a large proportion of MPZ lineage^+^ osteolineage cells in the frontal bone of newborn mice.(Chen et al., 2017) By contrast, our findings showed that MPZ lineage cells were largely restricted to nerve-lining SCs and adjacent iSCs with very few labeled osteolineage cells in the bone, regardless of age. This matches prior work with embryonic mice and adult fish.(Xie et al., 2019) From our observations, these findings suggest that SCPs are not a major source of skeletal lineages under homeostatic conditions. The presence of iSCs and potential for formation of repair SCs in adult bone nonetheless raises the possibility that this plasticity could be re-engaged under conditions such as injury or repair, warranting further investigation.

### Limitations and Additional Considerations

These findings provide important context for future studies aiming to genetically target SC populations in skeletal tissues. While MPZ-Cre robustly labels mature SC lineages, its constitutive developmental expression results in occasional labeling of non-glial derivatives. In contrast, p75-NGFR–based models preferentially label mature non-myelinating SCs and iSC-like populations but may also capture additional cell types throughout the body depending on context. Together, these observations indicate that no single marker fully captures the spectrum of SC lineages in bone, highlighting the need for careful experimental design, including inducible and combinatorial approaches.

It is also of note that in addition to single fibers, many nerves within the bone consist of multiple SC-covered axons running in parallel. This poses challenges for accurate quantification of total nerve axons and SCs in bone, particularly at low resolution. Previous studies have shown that nerve density in the periosteum of frontal and parietal bones is fairly similar to somewhat higher in the parietal bone.(Horenberg et al., 2025) However, we have found that the overall length density of Schwann cells is higher in the frontal bone than the parietal bone. This discrepancy could be due to the difficulty of imaging and quantifying very thin nerves. We have found the ZsGreen reporter to be extremely bright, allowing for quantification of areas of SCs covering nerves which were very faint with the TUBB3 antibody. At minimum, future studies are recommended to document the criteria for classification of a structure as “nerve”, and to provide detailed computational methods underlying any attempts to deconvolute nerves or SCs containing multiple vs single axons.

This study has several limitations that also define important directions for future work. While our analysis focuses on anatomical and cellular characterization rather than functional testing, it establishes a necessary framework for subsequent mechanistic studies. In addition, classification of mature SC and candidate iSC populations is based on marker expression and morphology, which may not fully capture the dynamic range of SC states. Technical aspects of imaging, especially in cleared tissues, may limit detection of SCs covering small axons and iSCs in complex whole tissue preparations. Finally, as our analyses were performed under homeostatic conditions, future studies will be needed to determine how these networks remodel in response to injury and disease.

## Conclusions

In conclusion, this work provides a foundational characterization of SC organization in the adult skeletal system. By demonstrating that SCs track with nerve networks, exhibit distinct morphological characteristics, and show region-specific distributions within bone, we position them as integral components of the bone microenvironment. These findings support a model in which SCs act at the interface of neural signaling and skeletal tissues, with relatively unexplored functional roles. Future studies integrating functional perturbation will be critical to define the mechanistic contributions of SCs to skeletal biology and disease.

## ACKNOWLDEGMENTS

This work was funded by grants from the National Institutes of Health (NIH) including R01-DK132073 (ELS) and R21-DE032420 (ELS). This work was also supported by the Musculoskeletal Research Center Cores and a Seed Grant from the Musculoskeletal Research Center (ELS) at Washington University (P30-AR074992), the WashU Genome Engineering and Stem Cell Core (GESC), microgrant funding from the Washington University Center of Cellular Imaging, and fellowship support from the Washington University Center of Regenerative Medicine (QM,MGH).

