## Supplemental Table and Figures for "Schwann Cell Mapping and Characterization in Bone of Different Embryonic Origins"

**Supplementary Table 1. Antibodies used for immunostaining of fixed tissue**

| **Primary Antibody (Vendor, Cat. No)** | **Storage** | **Dilution** | **Secondary Antibody** | **Fluorophore** | **Storage** | **Dilution** |
| --- | --- | --- | --- | --- | --- | --- |
| Anti-Tubulin Beta III  (Abcam, ab18207) | -20 ˚C | 1:1000 (calvaria)  1:500 (long bones) | Donkey Anti-Rabbit  (Jackson Immunoresearch, 711-605-152) | AlexaFluor-647 | -20 ˚C in 50% glycerol | 1:500 (calvaria)  1:300 (long bones) |
| Anti-S100B  (Millipore Sigma, HPA015768) | -20 ˚C | 1:1000 (calvaria) | Donkey Anti-Rabbit  (Jackson Immunoresearch, 711-605-152) | AlexaFluor-647 | -20 ˚C in 50% glycerol | 1:500 (calvaria) |
| Anti-p75 NGF Receptor  (Abcam, ab227509) | -20 ˚C | 1:1000 (calvaria) | Donkey Anti-Rabbit  (Jackson Immunoresearch, 711-605-152) | AlexaFluor-647 | -20 ˚C in 50% glycerol | 1:500 (calvaria) |
| Anti-GFP  (Abcam, ab13970) | -20 ˚C | 1:1000 (calvaria)  1:500 (long bones) | Donkey Anti-Chicken  (Jackson Immunoresearch, 703-545-155) | AlexaFluor-488 | -20 ˚C in 50% glycerol | 1:500 (calvaria)  1:300 (long bones) |

**SUPPLEMENTAL FIGURE 1**

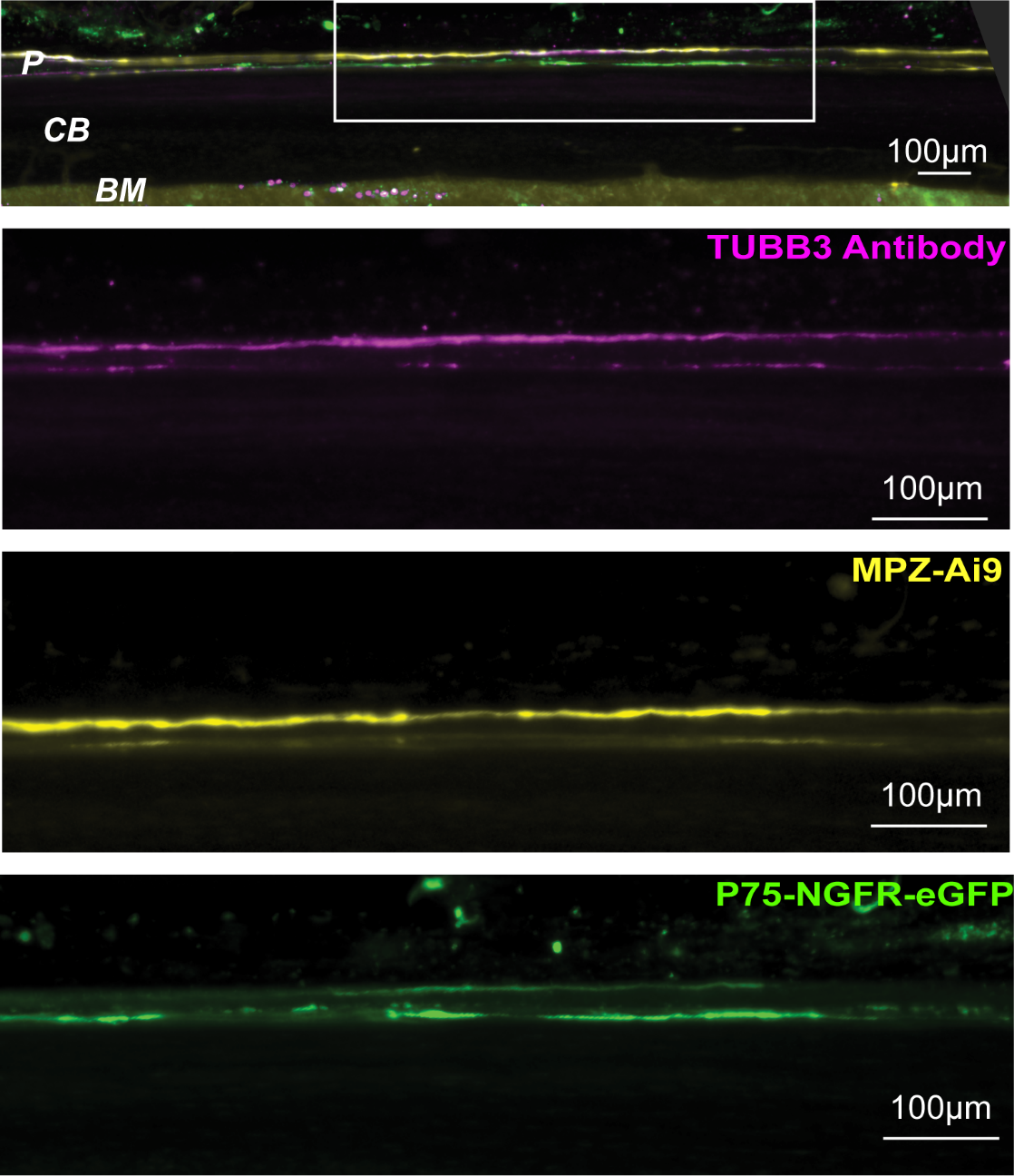

**Supplemental Figure 1.** **Schwann cells and nerves in the mouse tibial periosteum.**
Maximum intensity projection of whole mount 4x lightsheet microscopy images of ‘Schwann Cell Mapper’ (*Mpz*-Cre^+/-^;TdT^+/+^;*Ngfr*-eGFP^+/+^) mouse suggests a spatial distribution of the Schwann cells (SCs) within the periosteum, with myelinating MPZ-positive SCs localized to the outer periosteal layer and non-myelinating p75-NGFR-positive SCs enriched in the inner periosteal layer. Magenta = TUBB3 antibody, Yellow = MPZ-Ai9 reporter for myelinating SCs, Green = p75-eGFP reporter for non-myelinating SCs. Scale = 100 µm. P= Periosteum, CB= Cortical Bone, BM = Bone Marrow.

**SUPPLEMENTAL FIGURE 2**

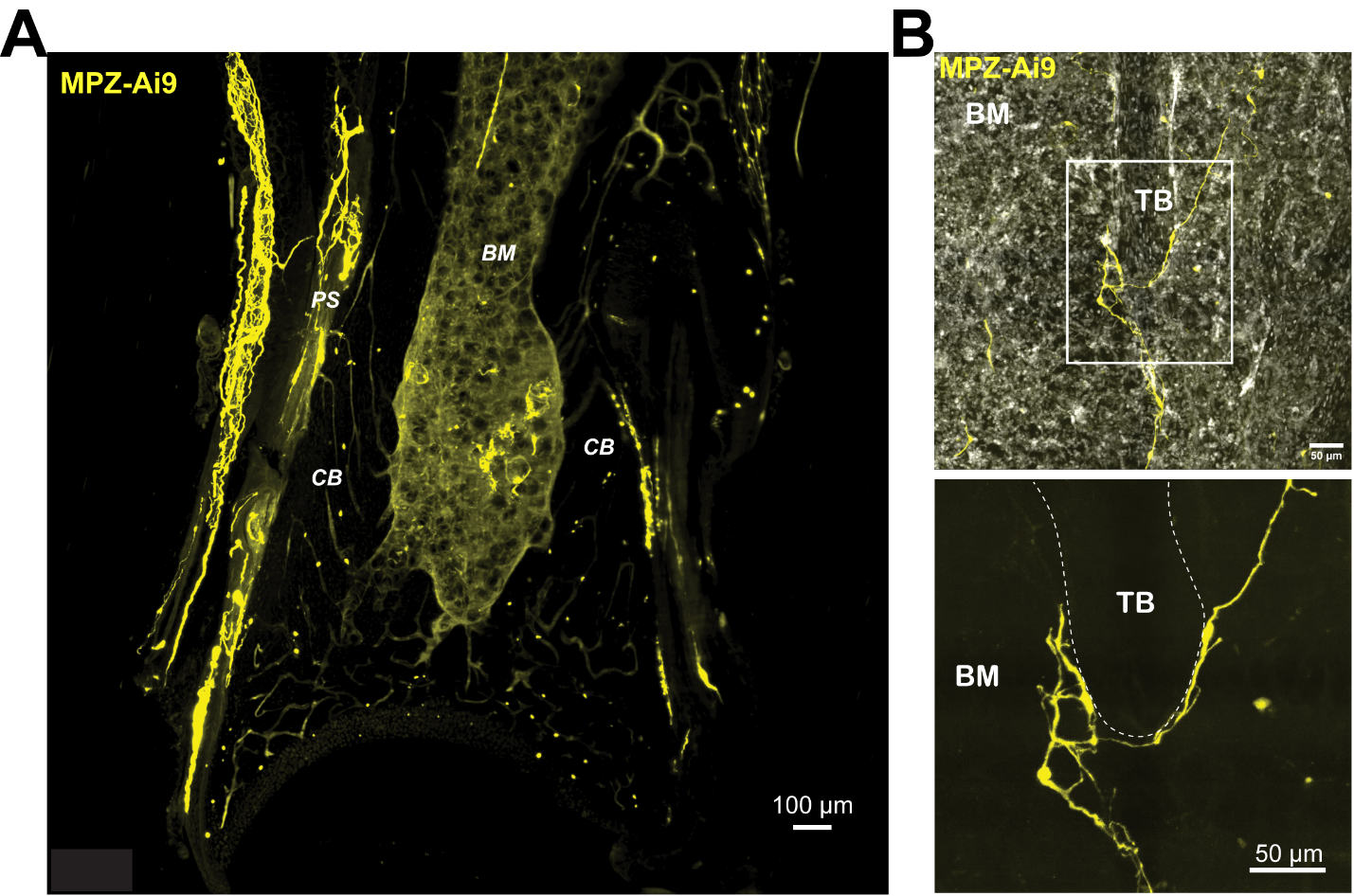

**Supplemental Fig 2.** **Schwann cells in periosteum and bone marrow space.** **(A)** Maximum intensity projection (MIP) of lightsheet microscopy images of ‘Schwann Cell Mapper’ (*Mpz*-Cre^+/-^;TdT^+/+^;*Ngfr*-eGFP^+/+^) mouse distal tibia showing a concentration of myelinating MPZ+ SCs in the periosteum. Scale = 100 µm. **(B)** MIP of lightsheet microscopy images of a ‘Schwann Cell Mapper’ mouse caudal vertebra showing presence of myelinating MPZ+ SCs around a trabecula in the bone marrow space. The white box indicates a zoomed in portion to the bottom. Scale = 50µm. Yellow= MPZ-Ai9 reporter for myelinating SCs, Gray=DAPI. CB= Cortical Bone, TB= Trabecular Bone, BM= Bone Marrow, PS= Periosteum.

**SUPPLEMENTAL FIGURE 3**

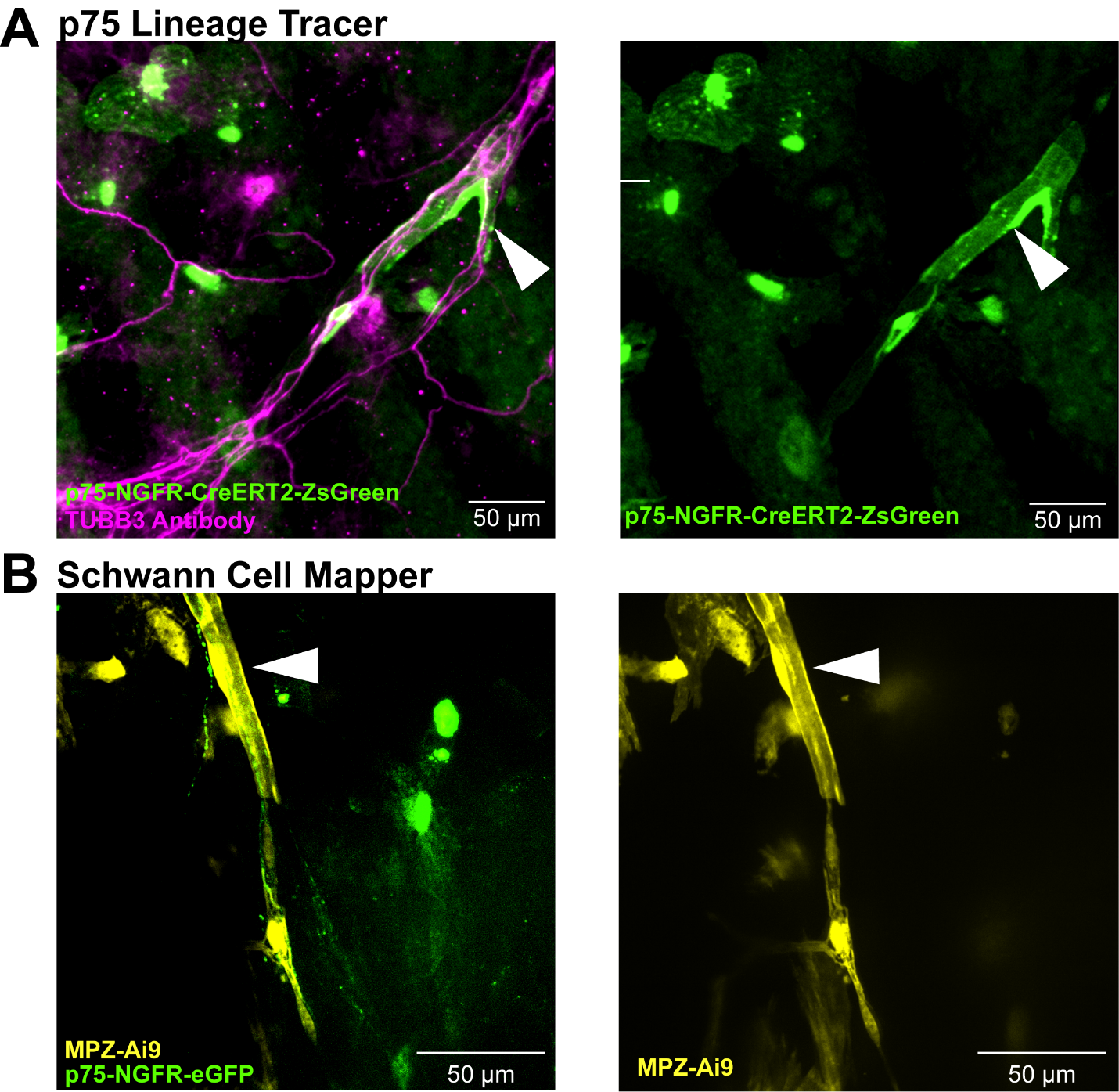

**Supplemental Figure 3. Immature Schwann Cells may transition into mature Schwann Cells. (A)** Maximum intensity projection (MIP) of whole mount 10x spinning disk confocal images of a ‘p75 Lineage Tracer’ (*Ngfr*-CreERT2^+/-^;ZsGreen1^+/+^) mouse calvaria showing a candidate p75-NGFR+ immature Schwann Cell (iSC) in transition. Magenta = TUBB3 antibody. Green= p75-NGFR-CreERT2-ZsGreen reporter. **(B)** MIP of whole mount 40x spinning disk confocal images of a ‘Schwann Cell Mapper’ (*Mpz*-Cre^+/-^;TdT^+/+^;*Ngfr*-eGFP^+/+^) mouse calvaria showing a candidate MPZ+ iSC in transition. Yellow = MPZ-Ai9 reporter. Green = p75-NGFR-eGFP reporter. Scale = 50µm.

**SUPPLEMENTAL FIGURE 4**

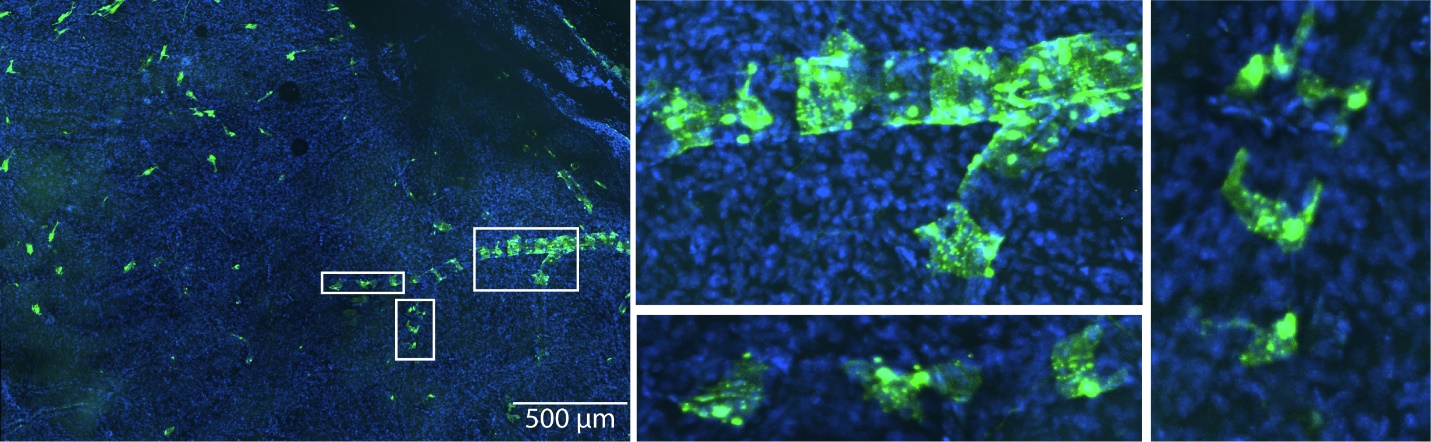

**Supplemental Figure 4. Additional p75-NGFR+ cells in calvaria.** 10x spinning disk confocal images of a ‘p75 Lineage Tracer’ (*Ngfr*-CreERT2^+/-^;ZsGreen1^+/+^) mouse calvaria showing p75-NGFR+ cells associated with a blood vessel-like structure. Green = p75-NGFR-eGFP reporter. Blue = DAPI.

**SUPPLEMENTAL FIGURE 5**

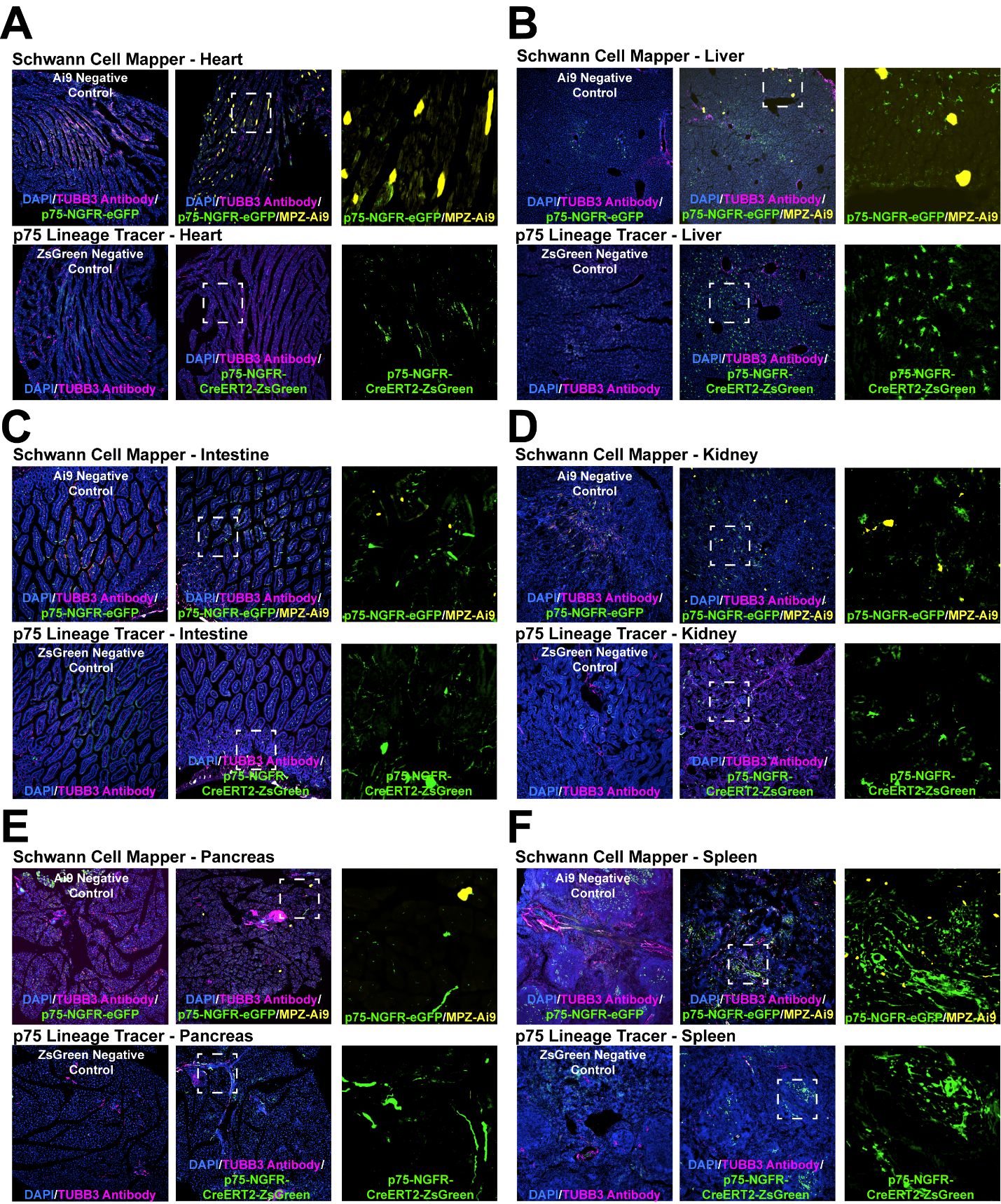

**Supplemental Figure 5. p75-NGFR+ and MPZ+ cells in various soft tissues.** Maximum intensity projection of 40x spinning disk confocal images of ‘Schwann Cell Mapper’ (*Mpz*-Cre^+/-^;TdT^+/+^;*Ngfr*-eGFP^+/+^) and p75 Lineage Tracer’ (*Ngfr*-CreERT2^+/-^;ZsGreen1^+/+^) mouse **(A)** heart sections, **(B)** liver sections, **(C)** intestine sections, **(D)** kidney sections, **(E)** pancreas sections, and **(F)** spleen sections, showing p75-NGFR+ and MPZ+ cells. Magenta = TUBB3 Antibody, Green = p75-NGFR-eGFP or p75-CReERT2-ZsGreen1 reporter, Yellow = MPZ-Ai9 reporter. Blue = DAPI.
